# Horizontal acquisition followed by expansion and diversification of toxin-related genes in deep-sea bivalve symbionts

**DOI:** 10.1101/605386

**Authors:** Lizbeth Sayavedra, Rebecca Ansorge, Maxim Rubin-Blum, Nikolaus Leisch, Nicole Dubilier, Jillian M. Petersen

## Abstract

Deep-sea bathymodioline mussels gain their nutrition from intracellular bacterial symbionts. Their sulfur-oxidizing (SOX) symbionts were recently shown to encode abundant toxin-related genes (TRGs) in their genomes, which may play a role in beneficial host-microbe interactions. Here, we compared TRGs in the genomes of SOX symbionts from 10 bathymodioline mussel and two sponge species to better understand their potential functions and evolutionary origins. Despite the close phylogenetic relatedness of these symbionts, the number and classes of encoded toxins varied greatly between host species. One of the TRG classes, YDs, has experienced gene expansions multiple times, suggesting that these genes are under adaptive selection. Some symbiont genomes contained secretion systems, which can play a role in host-microbe interactions. Both TRGs and secretion systems had a heterogeneous distribution, suggesting that these closely related bacteria have acquired different molecular mechanisms for interacting with the same family of animal hosts, possibly through convergent evolution.

## Introduction

Beneficial associations between animals and bacteria are virtually universal (McFall-Ngai et al. 2013). Many beneficial bacteria are acquired from the environment during host development, but the mechanisms that underpin host-symbiont recognition, invasion of host tissues or cells, and maintenance of the associations, are still not well understood (Pel and Pieterse 2013). In contrast, the molecular mechanisms pathogens use to interact with their hosts have been intensively studied (e.g. Sansonetti 2002; Di Genova and Tonelli 2016; Kaufmann and Dorhoi 2016; Kendall and Sperandio 2016). A number of pathogen-encoded proteins that interfere with host cell activity have been described and characterized as toxins (Lang et al. 2010; Aktories 2011; Huber et al. 2016). Large-scale bacterial genome sequencing has revealed toxin-related genes (TRGs) in the genomes of many beneficial bacteria with homology to characterized toxins of pathogens. This suggests that pathogens and beneficial bacteria use similar molecular mechanisms to interact with their hosts (Moya et al. 2008; Pérez-Brocal et al. 2011).

Bathymodioline mussels thrive at deep-sea hydrothermal vents and cold seeps by gaining nutrition from intracellular sulfur-and methane-oxidizing bacteria, which they harbor in their gill cells (Fisher et al. 1987; Duperron et al. 2009; Ponnudurai et al. 2016). Sayavedra et al. (2015) recently discovered diverse and abundant TRGs in the genomes of the sulfur-oxidizing (SOX) symbionts from two *Bathymodiolus* mussel species. These TRGs were hypothesized to play a role in beneficial host-microbe interactions, including host-symbiont communication and defense against parasites (Sayavedra et al., 2015). The sulfur-oxidizing (SOX) symbionts are acquired from the environment by each new host generation (Won et al. 2003; Wentrup et al. 2014), but little is known about the mechanisms the symbionts use to invade and survive within host cells.

In this study, we investigated the distribution of toxin-related genes (TRGs) in the SOX symbionts of ten *Bathymodiolus* species and in the closely-related SOX symbionts of two deep-sea sponge species. We hypothesized that TRGs encoded by all symbionts associated with a certain animal group (mussels or sponges) would be essential for interactions with their animal host such as recognition and invasion of host cells. Furthermore, given that TRGs were most likely acquired by the SOX symbionts through horizontal gene transfer, we aimed to understand how TRG acquisition has influenced genome evolution in this closely-related group of symbiotic bacteria.

## Results and Discussion

### Phylogenomic analyses reveal two well-supported symbiont clades

Previously, genome sequences were available from the SOX symbionts of three bathymodioline species from vents in the Pacific and Atlantic Oceans (Ikuta et al. 2015; Sayavedra et al. 2015). We sequenced and assembled the draft genomes of SOX symbionts from seven additional mussel species from vents and seeps around the world (Table 1, Supplementary Table 1). Furthermore, we assembled SOX symbiont genomes from metagenomes of three poecilosclerid sponges from the Gulf of Mexico that co-occur with two of the bathymodioline species investigated in this study (Rubin-Blum et al. 2017). The draft genomes sequenced in this study were between 90.8 to 98.5% complete, and were sequenced to depths ranging from 24x to 3600x. Their estimated genome sizes ranged from 1.41 to 2.82 Mbp. Many of these symbiont genomes may thus be larger than the only closed genome, that of the SOX symbiont of *B. septemdierum,* which is 1.47 Mbp (Ikuta et al. 2015).

**Table 1.**
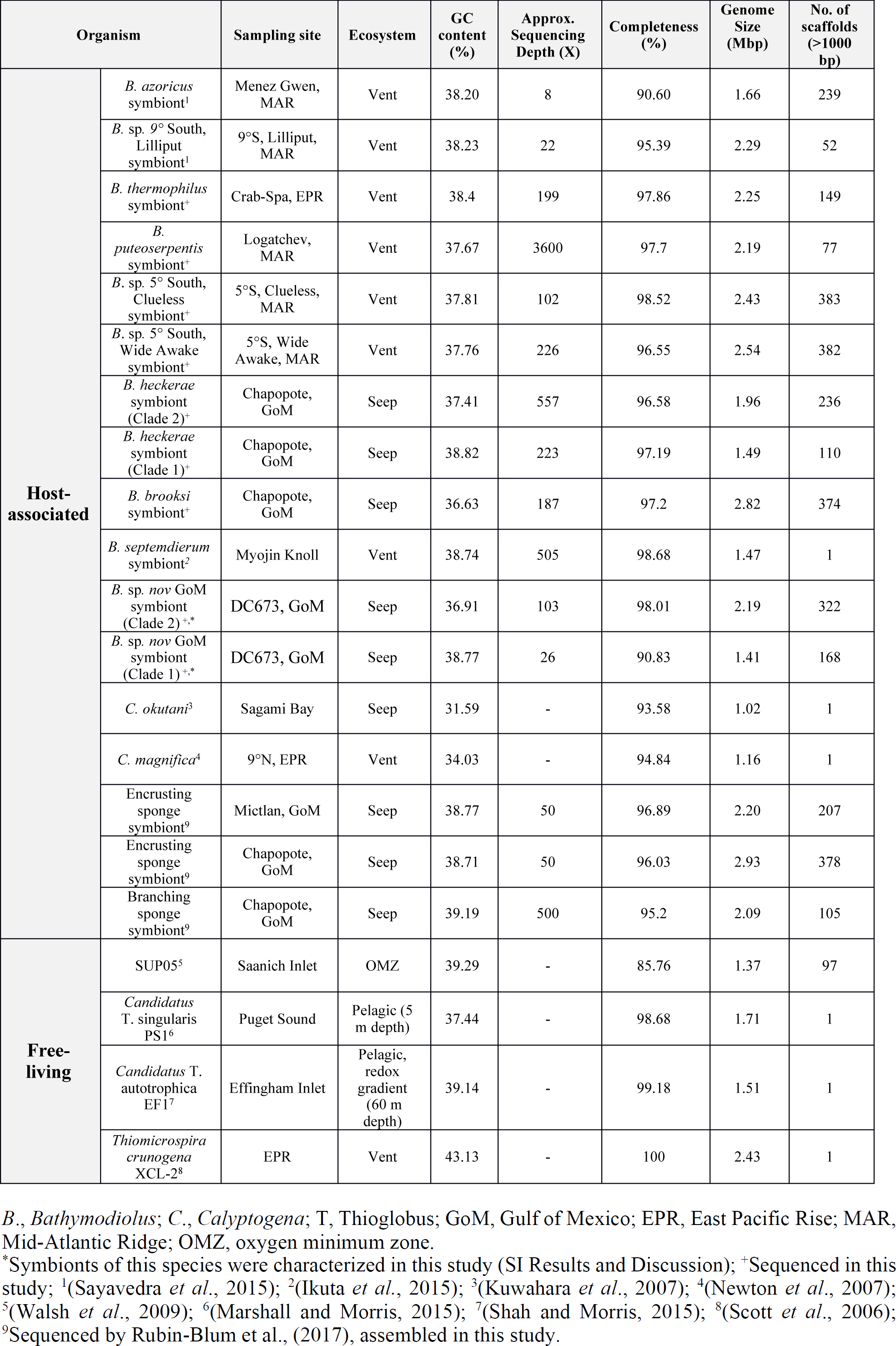
Overview of sulfur-oxidizing bacteria analyzed in this study.

We constructed a well-supported phylogenomic tree with 38 orthologous protein-coding genes from the sponge and mussel SOX symbionts and their close relatives. Consistent with previous 16S rRNA phylogenies, the SOX symbionts from mussels, sponges and clams did not form a monophyletic clade, as they were interspersed with free-living sulfur-oxidizing bacteria called ‘SUP05’ (Petersen et al. 2012; Sayavedra et al. 2015). The sponge-associated SOX symbionts formed a cluster together with most *Bathymodiolus* SOX symbionts, which we termed Clade 1 (Fig. 1 and Fig. 2). The symbionts of two mussel species, *B. heckerae* (BheckSOX) and *B*. sp. nov GoM (BspGoMSOX), clustered in a separate well-supported clade, together with the cultivated sulfur oxidizer *Candidatus* Thioglobus autotrophica EF1 and SUP05 bacteria from the Pacific Northwest (Clade 2, Fig. 1). The intermixing of symbiotic and free-living bacteria in our phylogenomic analysis, and in previous 16S rRNA phylogenies, suggests that either 1) free-living SOX bacteria acquired the ability to associate with bathymodioline mussels multiple times or 2) the free-living bacteria that fall within the highly supported clade of SOX symbionts from mussels, sponges and clams evolved from a symbiotic ancestor. So far, there is no evidence that these symbionts have a free-living stage that is metabolically active, although very closely-related free-living bacteria from the SUP05/*Ca. Thioglobus* clade are often abundant in hydrothermal vent environments (Anantharaman et al. 2012; Meier et al. 2017). In fact, the symbionts may rely on their hosts for some essential metabolites since they appear to lack two enzymes considered to be critical for anaplerotic metabolism (Ponnudurai et al. 2016). However, the isolate *Ca*. Thioglobus autotrophica also lacks one of these central metabolic enzymes: malate dehydrogenase. Thus, a free-living existence may be possible without enzymes previously assumed to be essential. We cannot rule out either of our two explanations above, but clearly, the well-supported clustering of sponge and mussel symbionts suggests that they shared a common ancestor, possibly undergoing a host-switching event, as well as multiple lifestyle switches from free-living to symbiotic and possibly symbiotic to free-living.

**Fig 1.**
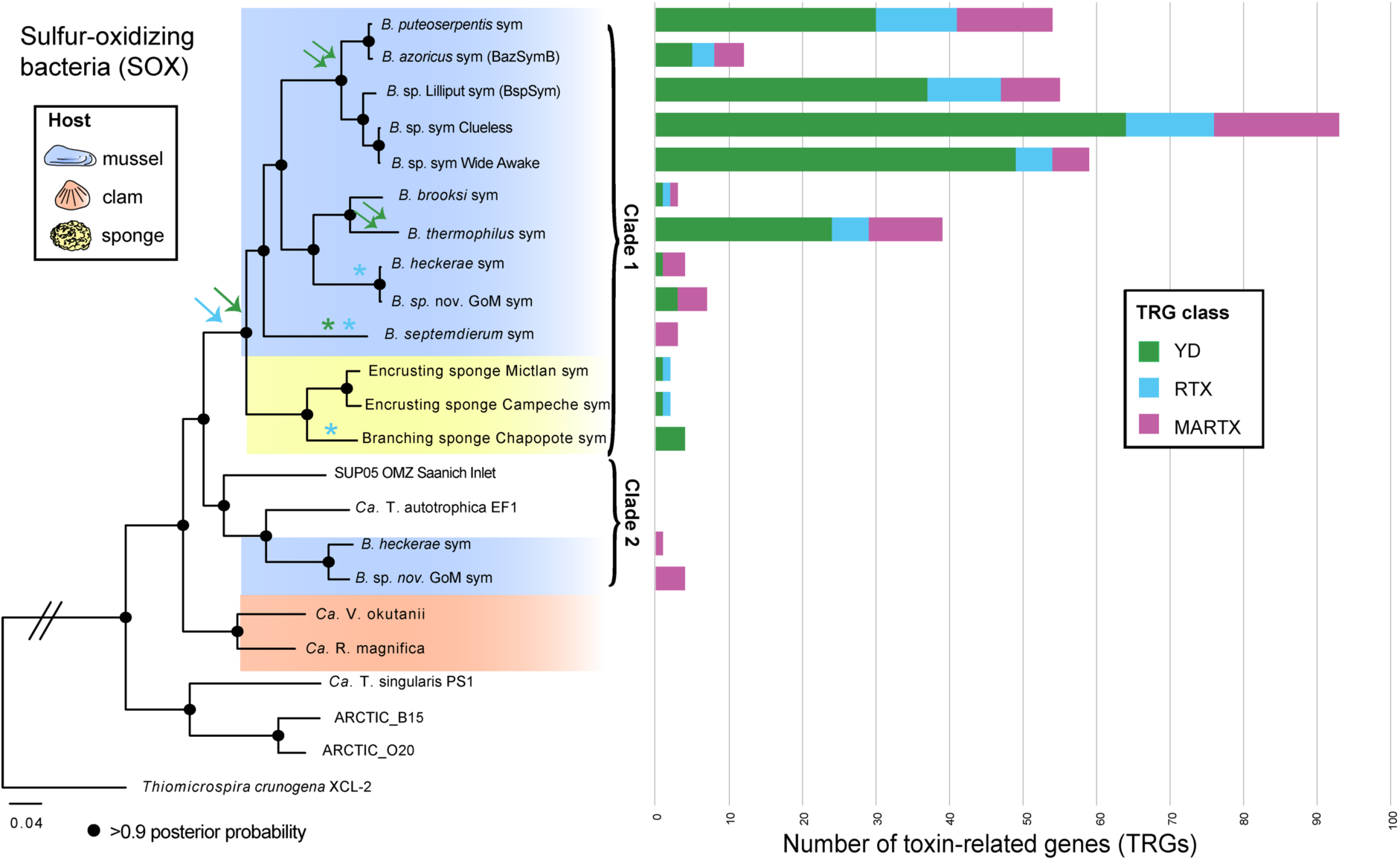
Phylogenomic tree of symbiotic and free-living SOX, estimated with 38 orthologous protein-coding genes, and the corresponding distribution of TRGs in these SOX. Filled circles represent a posterior probability higher than 0.9. The blue arrow indicates the proposed acquisition of RTX genes, green arrow of YD-repeat genes. All genomes shown in this tree were searched for all TRGs. Single arrows represent possible gene acquisition; double arrows indicate possible gene duplications; asterisks show possible gene loss events. The color of the arrows and stars corresponds to the TRG class. The number of individuals sequenced per species and geographic location is shown in Supplementary Table 1. T = *Ca*. Thioglobus; B = *Bathymodiolus*; sym = symbiont.

**Fig 2.**
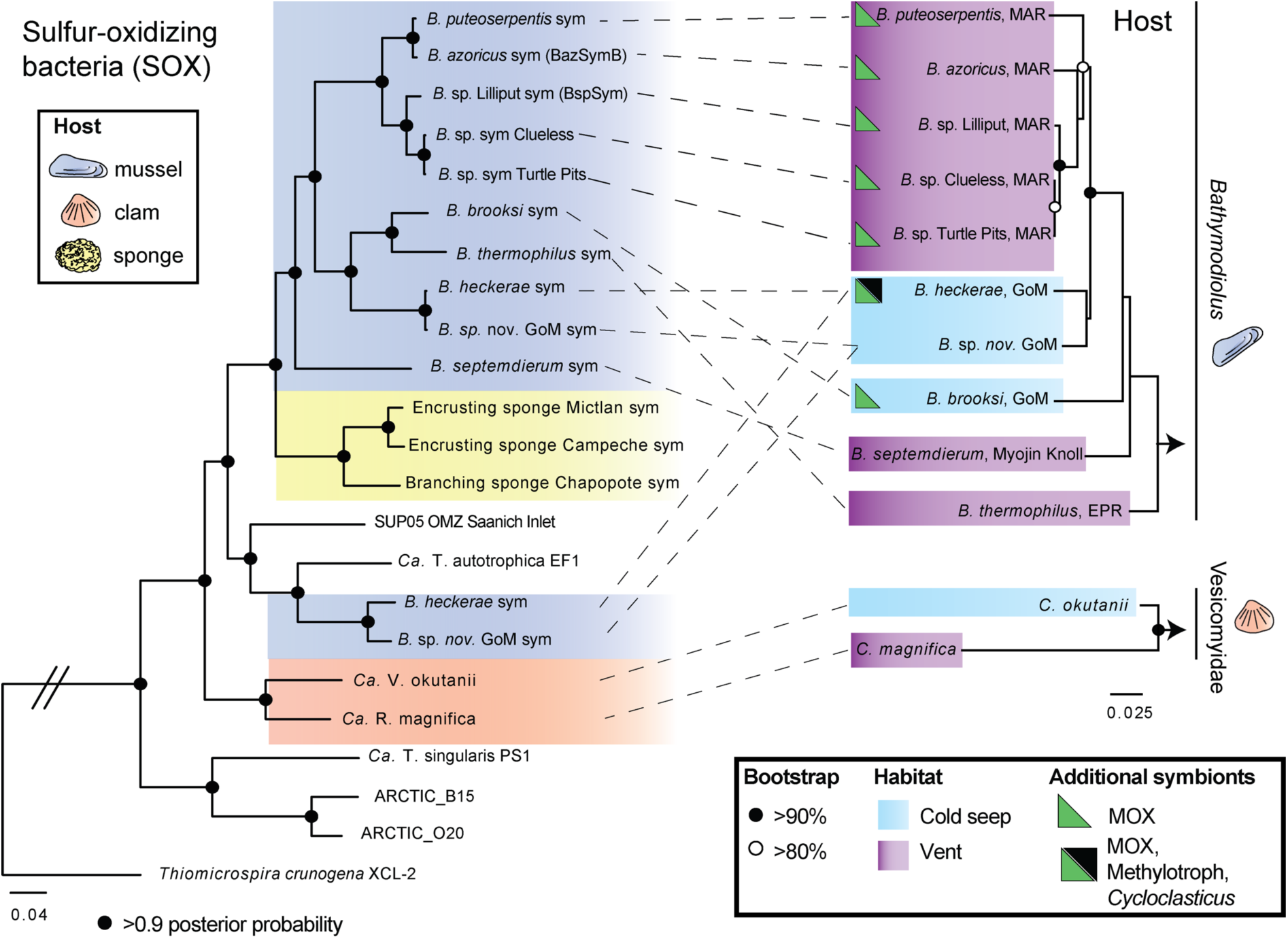
Bivalve phylogeny and symbiont phylogeny. The maximum-likelihood host phylogeny was reconstructed based on cytochrome oxidase I (COI). Symbiont phylogeny was estimated with 38 orthologous protein-coding genes.

### Horizontal acquisition, expansion and diversification of toxin-related gene families

SOX symbiont genomes from the two mussel species described by Sayavedra et al. (2015) encoded TRGs from three toxin classes: 1) RTX, or ‘repeat in toxin’ proteins, 2) MARTX toxins, which are large proteins containing multiple repeat motifs and domains of diverse functions, and 3) YD repeat toxins, named for their characteristic repeat sequence. In this study, we searched for TRGs in the SOX symbiont genomes of eight additional mussel species and two sponge species, as well as their closest free-living and symbiotic relatives (Table 1) (see SI Materials and Methods).

We consistently found TRGs in the SOX symbiont genomes of mussels and sponges, and these were highly abundant in the symbionts of mussel species (Fig. 1). In contrast, none of the genome sequences from bacteria closely related to the mussel and sponge SOX symbionts, such as free-living SUP05 and the vertically-transmitted, obligate intracellular symbionts of clams, encoded TRGs (Fig. 1).

#### MARTX

One toxin class, MARTX, was found in all of the mussel SOX symbiont genomes, regardless of whether they belonged to Clade 1 or 2. Intriguingly, MARTX were not found in any of the sponge symbiont genomes, even though these symbionts formed a highly-supported phylogenomic cluster together with mussel SOX symbionts. MARTX-like genes are known to be enriched in the genomes of symbiotic and pathogenic bacteria that associate with eukaryotes, and often have domains involved in attachment (Satchell 2011). The presence of MARTX-like genes in all mussel SOX symbionts from two distinct clades, and their absence in closely related free-living bacteria and the symbionts of clams and sponges, is consistent with a role in specific interactions with the mussel hosts, which could include attachment and recognition during colonization and intracellular infection of host gill cells. The length, sequences, domain content and arrangement of MARTX genes were highly diverse as shown previously for symbionts of two mussel species (Sayavedra et al. 2015) (Fig. 3). Despite this variable domain architecture, the symbionts of all 10 mussel species investigated had at least one MARTX-like gene with domains involved in attachment such as haemmagglutinin, cadherin, and integrin, indicating their role in attachment to host cells (Supplementary Table 2). If they are involved in attachment, they might also play a key role in mediating recognition and specificity. The mussel SOX symbioses are clearly highly specific: all except one of the known host species associate with only one or two 16S rRNA SOX types, which are not found in any other mussel species (see Duperron et al. 2008 for the only known exception). This host specificity is strictly maintained even when multiple mussel species co-occur, such as *B. brooksi* and *B. heckerae* at cold seeps in the Gulf of Mexico (Raggi et al. 2013). The highly divergent sequence and domain architectures of the MARTX genes in different symbiont lineages might be one of the mechanisms that determine this specificity. Although lacking MARTX genes, the SOX symbionts of sponges encoded proteins with leucine-rich repeats and cadherin domains, which have been hypothesized to play a role in recognition in shallow-water sponge symbioses (Thomas et al. 2010; Hentschel et al. 2012).

**Fig 3.**
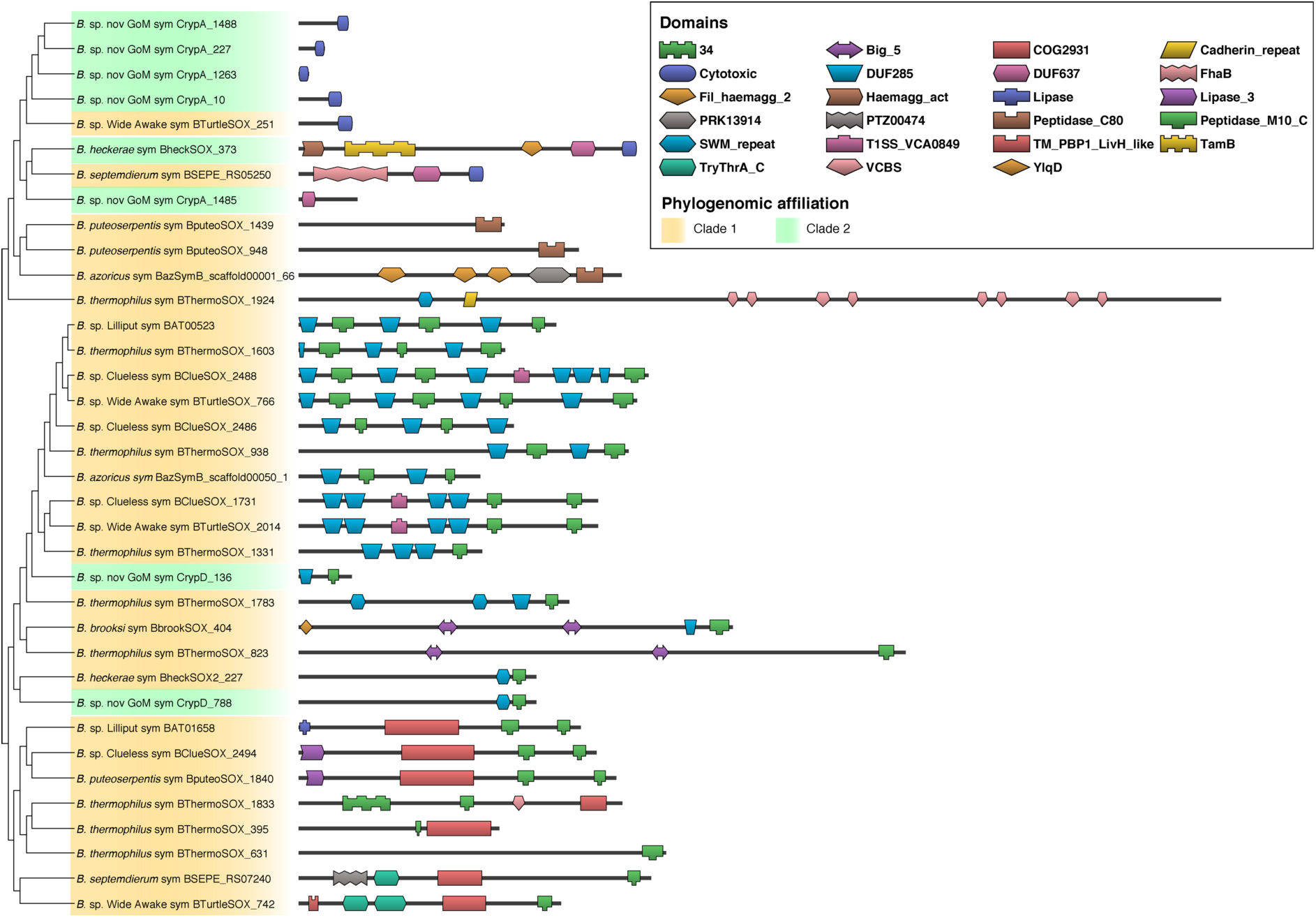
Domain structure of MARTX-like genes that have the most similar domain architecture to the MARTX-like genes from Clade 2 SOX symbionts (Fig. 1). The tree was estimated with the domain distance among proteins with DoMosaics (Moore et al. 2014). The descriptions of these domains are available in Supplementary Table 2. Locus tags of the MARTX genes from the SOX symbionts are shown at the nodes.

#### RTX and YD repeats

The second toxin class, RTX, was found in some but not all mussel and sponge symbionts from Clade 1, and not in any of the Clade 2 symbionts. The third class of genes, YDs, was found in all members of Clade 1 except the basal *B. septemdierum* symbiont (Fig. 1). Clade 2 symbionts did not contain any YD repeat genes, but these symbionts co-exist in a dual symbiosis with Clade 1 symbionts that did encode YD repeats (Fig. S1). These observations support the following hypotheses: I) RTX and YD repeats are not essential for establishing and maintaining an intracellular symbiotic association with mussels, II) RTX genes were acquired by the common ancestor of Clade 1 and lost on multiple occasions, III) YD genes were acquired by the common ancestor of Clade 1, and YD genes were subsequently lost in the *B. septemdierum* symbiont, and IV) gene duplication contributed to the expansion of the YD genes (SI Results and Discussion). Given that YD and RTX appear to not be essential for intracellular symbiosis, their main role might be to defend their hosts against possible pathogens or parasites (SI Results and Discussion).

#### Secretion system genes

Secretion systems (SS) are often essential for pathogens to survive inside host cells (Green and Mecsas 2016). We therefore searched for SS components in the genomes of the SOX symbionts and their free-living relatives. We found genes encoding components of almost all known SS types. Like the TRGs, these SS components were patchily distributed among the SOX symbiont genomes, and not a single SS was specific to all of the intracellular bacteria (Supplementary Table 3).

All genome bins from SOX symbionts of Clades 1 and 2 encoded VgrG, a component of the type VI SS (T6SS). Although none of the genomes analyzed in this study encoded the full suite of T6SS genes, VrgG alone may allow the export of toxins without the full T6SS gene array (Hachani et al. 2014). This gene was also present in some of the free-living SOX relatives. Three genes considered essential for T4SS were present in three of the thirteen SOX symbionts of Clade 1 and in both SOX symbionts of Clade 2, but not in any of the free-living or clam SOX. Most of the T4SS present in the mussel SOX symbionts encoded a relaxase that can interact with DNA, supporting a role in conjugation (Abby et al. 2016). In some pathogens, the same T4SS can carry out dual functions in conjugation and host colonization (Dehio 2008).

The BheckSOX of Clade 2 encoded an additional T4SS of the type VirB/D, which was not found in any other mussel SOX symbionts. The BheckSOX VirB/D-T4SS shares a similar genomic architecture with systems used for both conjugation (e.g. *Vibrio parahaemolyticus*), and for host cell invasion and persistence through secretion of toxic effectors (e.g. *Bartonella henselae* str. Houston-1) (Seubert et al. 2003; Schmid et al. 2004; Dehio 2008; Gokulan et al. 2013). A phage integrase was found upstream of the VirB/D T4SS gene cluster, raising the possibility that, just as in pathogens, beneficial bacteria could be acquiring and exchanging secretion systems from bacteriophages (Guy et al. 2013).

## Conclusions

The SOX symbionts of deep-sea mussels and sponges encoded a highly diverse array of toxin-related and secretion system genes. Our comparative genomic analyses identified only one toxin class, MARTX, which was common to all mussel SOX symbionts and might therefore be a gene class essential for host-microbe interactions such as recognition, attachment and symbiont uptake in the mussel symbioses. All other TRGs and secretion systems had a heterogeneous distribution in the symbionts we investigated, which attests to the complex and varied routes of genome evolution taken by the members of this closely-related group of symbiotic bacteria. If the SOX symbionts use their species-specific sets of TRGs and secretion systems to interact with their respective hosts, this would be an example of convergent evolution in which free-living bacteria took multiple unique evolutionary trajectories to become intracellular symbionts of animals, depending on the genes they acquired.

TRGs and T4SS that could export protein effectors were not present in free-living SUP05, even though these bacteria are often found in hydrothermal vent plumes in close proximity to mussels (Sylvan et al. 2012; Anantharaman et al. 2014). It is therefore likely that these genes were acquired from other free-living or host-associated bacteria. Gene flow between these bacterial donors, SUP05 bacteria, and SOX symbionts in a ‘free-living’ stage in the environment could lead to the evolution of novel symbiont and free-living lineages (SI Results and Discussion) (Roux et al. 2014, our own unpublished data). Further investigation of horizontal gene transfer and genome evolution in groups of closely related bacteria such as the SUP05 and SOX symbionts, could reveal how free-living bacteria become symbionts.

Some pathogen groups such as *Pseudomonas aeruginosa* show a similar pattern to the SOX symbionts we investigated, with species-or strain-specific differences in their genomic complement of toxins and virulence factors (Huber et al. 2016). In *P. aeruginosa*, these genomic differences are clearly reflected in major phenotypic differences such as severity of human disease. At the morphological level, the SOX symbionts of different mussel and sponge species do not show clear differences. However, just as in pathogens, the underlying genomic variation between symbionts could result in differences in the way these diverse symbionts interact with their hosts. For example, some host species seem to consistently carry a higher symbiont load than others (Duperron et al. 2008; Raggi et al. 2013), and this could not only be due to differences in the availability of their energy sources, but also to differences in the rates of symbiont acquisition, maintenance, proliferation and digestion by the host. TRGs and SSs are likely to affect such host-microbe interactions and could thus have a significant impact on the functioning and stability of these symbioses (Huber et al. 2016).

## Acknowledgements

We thank the captains, crews and funding agencies of the sampling cruises AT26-23, M64-2, M67-2, M78-2, ATA57, M114-2 and NA043. We thank Christian Borowski, Stephanie Markert, and Charles Fisher for providing samples, Brandon Seah for helpful discussions and Miriam Sadowski for technical assistance. This work was funded by the Max Planck Society, the DFG Cluster of Excellence “The Ocean in the Earth System” at MARUM (University of Bremen), a European Research Council Advanced Grant (BathyBiome, Grant 340535) and a Gordon and Betty Moore Foundation Marine Microbiology Initiative Investigator Award through Grant GBMF3811 to ND, the DAAD through a doctoral grant to LS, and the Vienna Science and Technology Fund (WWTF) through project VRG14-021 to JMP.

## Author contributions

LS, ND, and JP conceived the study; LS, RA, and MRB analyzed the data; NL did TEM analysis; LS and JP wrote the paper with contributions and revisions from all coauthors.

