## Supplementary information for "Horizontal acquisition followed by expansion and diversification of toxin-related genes in deep-sea bivalve symbionts"

### SI Results and discussion

#### Were YD repeat genes acquired through multiple gene duplication or independent HGT events?

In genomes containing YD repeat genes, we always found multiple distinct copies. This could be the result of gene duplication, or multiple independent HGT events. If the YDs were the result of independent HGT events, the different gene copies found in a single genome would be expected to have closely-related homologs in other bacterial taxa (Kuo and Ochman 2009). In contrast, if multiple YDs are the product of gene duplication, we would expect that these YDs within a symbiont genome are most closely related to each other and thus cluster together in phylogenetic analysis (>30% amino acid similarity and >60% coverage) (Domman et al. 2014). Our results suggest that gene duplications were the main factor leading to multiple gene copies: firstly, based on phylogeny, the YDs from the SOX symbionts were distinct from all other sequences in public databases and formed two separate subclusters, A and B (>80% bootstrap support) (Fig. S1). Secondly, species-specific expansion events were observed in the genomes of the SOX symbiont of *B. thermophilus* (BthermoSOX), as well as the SOX symbionts from the Southern Mid-Atlantic Ridge (SMAR) (the SMAR symbionts belong to a single bacterial species based on an average nucleotide identity between 96.6 to 99.7%). Thirdly, the multiple copies of YDs encoded within single symbiont genomes were most similar to each other (Fig. S1). Finally, the YD genes were often in tandem order. These observations strongly suggest that tandem gene duplication contributed to the expansion of the YD genes in some of the mussel SOX symbionts.

#### Are YD and RTX genes involved in defense?

Considering that YD and RTX genes do not seem essential for intracellular symbiosis with mussels because the symbionts of some hosts do not have them, could their main function be protecting the host against pathogens and parasites (Sayavedra *et al.* 2015)? The selective advantage for a symbiont that can protect its host would depend on multiple factors including host immunity and the abundance of parasites and pathogens in the host's environment (Hillman and Goodrich-Blair 2016). The expansion of the YD repeat gene family in Clade 1 symbionts is consistent with a role in defense, since a diverse range of toxins would be advantageous against counter-defense mechanisms of the parasite (Nuismer and Otto 2004; Tellier et al. 2014). If the YD repeat proteins are involved in defense, then the mussels that host YD-encoding symbionts may either experience more intense pressure from parasites and pathogens in the environment, or their hosts might have lost certain immune factors over evolution. For example, the marine invertebrates *Nematostella* and *Hydra* have lost part of the complement effector pathway (Miller et al. 2007). Hosts with defensive symbionts can also become 'addicted' to symbiont defense, as selective pressure to maintain their own immunity is lowered in the presence of the defensive symbiont (Martinez et al. 2016). The expansion of YDs in the symbionts of Clade 1 may have been accompanied by a loss of immune gene families in their respective mussel hosts. No complete genomes are available for *Bathymodiolus* that host SOX symbionts (Sun et al. 2017), it is therefore currently unknown if those mussel species that host YD-encoding symbionts have indeed lost part of their immune pathways. *B. thermophilus* and the *B. puteoserpentis/B.azoricus/B.*sp. species complex from the Mid-Atlantic Ridge would be ideal lineages to test this hypothesis, as their symbionts encode the most YD repeat genes. Like the YD repeats, RTX proteins were only found in some members of Clade 1 (Fig. 1). This could indicate that the RTXs also play a role in defense rather than essential host-symbiont crosstalk.

#### Origin of MARTX genes

To investigate whether the MARTX genes from Clade 1 and 2 mussel symbionts had a common origin, or whether each symbiont lineage acquired MARTX genes independently from different sources, we compared the sequence and domain architecture of the SOX symbiont MARTX genes with those in public databases. The genes from Clade 2 symbionts were most similar to each other (96.5% similarity, 48% coverage), and were more similar to genes from Clade 1 mussel symbionts (up to 51.5% similarity, 52% coverage), than to any other sequence available in public databases. Although we cannot rule out that a more similar MARTX gene exists in nature and is yet to be sequenced, the similarity between MARTX genes from the two distinct clades suggests a common source or origin. This can be explained by three scenarios: 1) the last common ancestor of Clades 1 and 2 had MARTX genes, and the free-living bacteria in Clade 2 and sponge symbionts in Clade 1 subsequently lost these genes, 2) bacteria we have not yet discovered were a source of genetic material that was transferred to the mussel symbionts in Clade 1 and 2 via HGT in two independent events, or 3) one of the mussel symbiont lineages acquired the MARTX and it was subsequently transferred to free-living close relatives we have not discovered yet, which evolved to become symbionts within the second clade. Mussel symbionts and free-living SUP05 bacteria co-occur at hydrothermal vents, and possible carriers of genetic material including viruses, outer membrane vesicles (OMV) and conjugation systems have been found in free-living and symbiotic SUP05 bacteria (Anantharaman et al. 2012; Roux et al. 2014; Meier et al. 2017) (see below for conjugation systems; Fig. S3 OMV). Although we cannot clearly identify which of the three scenarios is most likely based on the current data, our results raise the possibility that free-living and symbiotic sulfur oxidizers affiliated with the SUP05 clade may be genetically linked through HGT. TRGs may be shuffled between symbiont lineages or between symbionts and close relatives in the environment.

### Symbionts of *B*. sp nov Gulf of Mexico

The intracellular location of the bathymodioline SOX symbionts was confirmed in all species analyzed in this study, with the exception of the recently described *B*. sp. nov GoM (Faure et al. 2015). Based on the SSU rRNA, the two metagenomes of *B*. sp. nov GoM had the two SOX symbionts described in this study, as well as epibiotic epsilonproteobacteria (Assié et al. 2016), but not the methane-oxidizing symbiont (Fig. 2 and Fig. S3). Our TEM analyses revealed that the SOX symbionts of *B*. sp. nov GoM occurred inside host vacuoles in specialized host cells (bacteriocytes). The symbionts were found near the surface of the cell, in the apical region of the bacteriocytes (Fig. S3), as described for other bathymodioline mussels (Nelson et al. 1995).

### SI Materials and methods

#### Sample collection and DNA extraction

Sampling sites and coordinates of the mussels and sponges sequenced and/or assembled in this study are listed in Supplementary Table 1. The gills of *B.* sp from 5° SMAR (Wide Awake) were dissected and homogenized in 1x PBS within a few hours of sampling. The homogenate was immediately centrifuged at 100 g for 10 min and the supernatant filtered through a series of GTTP filters: 12 µm, 5 µm, and 2 µm. The 2 µm cellulose acetate filter (Millipore, Germany) had an enriched fraction of SOX symbiont cells and was stored at -80°C. All other mussel species were dissected on board after collection and their gill tissues stored directly at -80°C (*B. puteoserpentis, B.* sp from 5° SMAR (Clueless), *B. brooksi, B. heckerae*, *B*. sp. nov. GoM), or first fixed in 96% ethanol and then stored at -80°C *B. thermophilus*), or fixed in Trump's fixative (McDowell and Trump 1976) for transmission electron microscopy (TEM) and stored at 4°C. In the home laboratory, gill tissues were transferred to innuSpeed Lysis tubes E (Analytic Jena, Germany) and homogenized in a FastPrep-24 Instrument (MP Biomedicals, Germany) for 2 min at 4 m/s. DNA was extracted from the homogenates and the cellulose acetate filter with cells as described by Zhou *et al*. (1996) or with the DNeasy blood and tissue kit (Qiagen, Germany) (Supplementary Table 1). Sample collection and DNA extraction of the sponges is described in detail by Rubin-Blum et. al., (2017). Shortly, the sponges were fixed in RNAlater (Sigma, Germany) on board. In the home laboratory DNA was extracted with the AllPrep DNA/RNA MiniKit (Qiagen, Germany).

#### Sequencing, assembly and genome annotation

Genomic DNA libraries were generated with the Illumina TruSeq DNA Sample Prep Kit (BioLABS, Germany), one library each for between one and three individuals per species or sampling location. Sequencing details are shown in Supplementary Table 1. Adaptors were removed and the reads were quality filtered (Q=2) with BBDuk V35 (Bushnell B. - sourceforge.net/projects/bbmap/). One initial assembly was done for the metagenomic reads of all individuals sequenced for each geographic region or mussel species with IDBA-UD V1.1.1 (Peng *et al.*, 2012). Sponge metagenomes (Rubin-Blum et al. 2017) were assembled for each host individual. The metagenome assemblies were binned with gbtools (Seah and Gruber-Vodicka 2015) based on differential genome coverage, taxonomic affiliation and GC content as described by Albertsen *et al.* (2013). The resulting genome bin was used as a reference to map the reads from each host individual from the same site or species, if more than one was sequenced, with BBMap V35 using a similarity cutoff of 98%. The reads from each individual mussel or sponge that mapped to this reference genome bin were used for a separate re-assembly with Spades V. 3.6, to improve the assembly results. This resulted in high-quality draft genomes for each individual that were sequenced when the genome coverage was at least higher than 10x (Supplementary Table 4). These individual assemblies were re-binned to remove possible contaminant contigs with gbtools based on differential genome coverage, taxonomic affiliation and GC content. In comparisons between sites and species where multiple individuals were sequenced, we only included the draft genome from the individual with the highest SOX symbiont coverage in Fig. 1, but the analysis of TRG distribution was done on all draft genomes (Supplementary Table 4). Contigs smaller than 800 bp were discarded from the final bins (Bouck et al. 1998; Luo et al. 2012). The assembly of *B. puteoserpentis* SOX was gap-filled with GapFiller (Boetzer and Pirovano 2012) since assembly of mate-paired sequencing reads from this sample introduced large stretches of Ns. Completeness was estimated with the proteobacterial UID3880 marker set from CheckM (Parks *et al.*, 2015). Average nucleotide identity (ANI) was estimated with the enveomics collection (Rodriguez-R and Konstantinidis 2016).

#### Investigating the distribution of TRGs and SS in symbiont genomes

Fragmented genomes pose a challenge for comparative genomics. To provide additional support for the observation that RTX and YD genes are absent in the SOX symbiont genomes of *B. heckerae and B.* sp nov, we scanned the entire metagenome assembly produced by IDBA-UD, including contigs smaller than 800 bp (Peng et al. 2012), with Diamond (Buchfink et al. 2014) using the collection of RTX and YD protein sequences of the SOX symbionts of Clade 1 as 'bait'. Although using a known sequence as bait might not include highly divergent sequences, we assumed that the proteins among closely related symbionts would be detectable using sequence similarity methods. Some contigs in the SOX symbiont of *B. heckerae* and *B*. sp nov metagenomes showed sequence similarity to the YD genes in Cluster B (similarity between 32 – 40%, coverage between 51 – 88%). The genes that showed hits against TRGs were first annotated with the following approach: the proteins of the whole metagenome were predicted with Prokka (Seemann 2014). Protein sequences that had similarity to TRGs were functionally annotated based on the domains described below. Some contigs of *B. heckerae* and *B*. sp. nov GoM had YDs with the domains considered in this study. To confirm that these contigs did not belong to the SOX symbiont from Clade 2, we reassembled the metagenomes with metaSpades V. 3.8, which outputs an assembly graph (Nurk et al. 2016). This graph was visualized with Bandage (Wick et al. 2015) to look for potential connections between Clade 2 SOX symbiont genomes and contigs with genes that had YDs. The assembly graph did not connect these contigs, and the YD-containing contigs had a sequencing coverage that matched those of the SOX symbionts from Clade 1 (Fig. S2). Thus, we consider it more likely that these contigs belonged to the low-abundance SOX symbionts of Clade 1 that co-occur with the Clade 2 SOX symbionts in *B. heckerae* and *B*. sp. nov GoM (SI Results and Discussion). The same approach was used to confirm that the SOX symbionts of sponges did not encode MARTX, but we used the MARTX sequences as 'bait'. MARTX were not found in any of the sponge metagenome assemblies. Although closed genomes would provide conclusive evidence, we consider it likely that at least some of this patchiness truly reflects the absence of these genes, as we never found BLAST hits in the corresponding whole metagenomes.

The genomes were scanned for secretion systems with the models from TXSScan (Abby et al. 2016) incorporated into MacSyFinder (Abby et al. 2014). SS are encoded by operons containing many genes. So in those cases where whole operons were not found in symbiont genomes, we can be more confident that they are not present in these symbionts. Similar genome architectures of the T4SS of type VirB/D from the Clade 2 SOX symbionts were searched with RAST and IMG (Aziz et al. 2008; Markowitz et al. 2011). We confirmed that the VirB/D-T4SS was only present in BheckSOX as described above for the TRGs with the following modification; as a query, we used the VirB/D-T4SS system from BheckSOX against the entire metagenome assemblies of the mussel samples with symbionts from Clade 1. To confirm that the SOX symbiont of *B. septemdierum* did not encode secretion systems in plasmids, we re-assembled the published metagenome with the accession number: DRA002953. No hits were found in any of the whole genome assemblies with a BLAST e-value cutoff of 0.01. Based on the high read coverage of the SOX symbionts in all mussel individuals, and the lack of BLAST hits in metagenomes from multiple host individuals, it is highly likely that the VirB/D-T4SS is specific to BheckSOX.

#### Phylogenetic analyses and identification of toxin-related genes

The draft genomes described in this study were annotated with RAST (Aziz *et al.*, 2008). The shared orthologous sequences were obtained with OrthoMCL (Li 2003) implemented in Odose (Galaxy). The 38 trimmed aligned sequences were concatenated and positions that were missing in more than 90% of the sequences were removed with Geneious V. 9 (Kearse et al. 2012) resulting in 38,032 nucleotide positions. A phylogenomic tree was reconstructed with MrBayes v. 3.2.6 (Ronquist and Huelsenbeck 2003) using two heated chains and 300,000 generations.

Candidate TRGs were obtained with BLAST searches using the protein sequences of the TRGs described previously (Sayavedra *et al.*, 2015). We considered only those sequences that had a similarity higher than 25% and minimum query coverage of 25% or a minimum alignment of 200 amino acids. TRGs of the mussel SOX symbionts are polymorphic systems – the domains that are present within the genes often vary in presence and order. The annotation of RTX and MARTX-like genes can therefore be challenging. To overcome the limitations of automatic annotation, we designed a scoring script that predicts the class of the TRG based on the domain combination present in the protein sequence and protein length. One limitation of the domain scoring system is that genes without a detectable domain are not classified, but it allows a consistent and comparable method for identifying functional domains present in the TRGs. The protein sequences were submitted to the batch conserved domain database (CDD) website (Marchler-Bauer et al. 2014). The resulting file was used as input for a self-written Perl script that assigned a score to the domains of each protein based on how often the domain was present in the classes YD, RTX or MARTX (see Supplementary Table 5 for the scores assigned to the domains observed in TRGs). Only the proteins that had at least one functional domain were further considered (Fig. 1). If the protein class obtained using the domain score system differed from the class identified via BLAST, then the annotation was checked manually. The presence of peptidase 80 or multiple functional domains was considered to be characteristic for MARTX proteins.

To find proteins with similar domain content and structure to the MARTX protein of the BheckSOX, we reconstructed a domain distance tree with the domains obtained with CDD using DoMosaics (Moore et al. 2014). Overlapping domains were resolved based on their e-value. Symbiont proteins that had similar domains to the MARTX protein of BheckSOX are shown in Fig. S3.

To investigate YD gene expansions we constructed a phylogeny with the YDs from SOX symbionts and the most closely related sequences in the databases. YD protein sequences were aligned with MAFFT (Katoh et al. 2002). Positions that had more than 90% gaps were not further considered. The resulting alignment of 649 positions was used to reconstruct a maximum likelihood phylogeny with RaxML (Stamatakis 2006) using 100 bootstrap replicates. The host phylogeny of *Bathymodiolus* mussels and the vesicomyid clams was estimated with the mitochondrial cytochrome c oxidase 1 (CO1) (Fig. 2). CO1 nucleotide sequences were aligned with MAFFT (Katoh et al. 2002), producing an alignment of 1,171 positions. A phylogeny was reconstructed as described for the YD genes.

#### Transmission electron microscopy (TEM)

To show the intracellular location of symbionts in *B*. sp. nov GoM, we examined two mussel individuals with TEM. Before embedding, mussel gill pieces were fixed in 2.5% glutaraldehyde in marPHEM (1.5x PHEM containing 9% sucrose (Montanaro et al. 2016)) for 1 hour at room temperature, washed three times with marPHEM, post-fixed with 1% osmium tetroxide in marPHEM for 1 hour at 4°C and dehydrated using a graded ethanol series (30%, 50%, 70%, 80%, 90%, 100% twice) on ice. The samples were transferred into 100% dry acetone and infiltrated using centrifugation (modified from McDonald 2014) in 2 ml tubes sequentially with 25%, 50%, 75% and 2x 100% Agar Low Viscosity resin (Agar Scientific, Stansted, Essex, United Kingdom). For this infiltration process, the samples were placed on top of the liquid and centrifuged for 30 s with a bench top centrifuge (Heathrow Scientific, USA) at 2,000 g for each step. After the second pure resin step, they were transferred into fresh resin in embedding molds and polymerized at 60°C for 12 h.

Ultra-thin (70 nm) sections were cut with an Ultracut UC7 (Leica Microsystem, Vienna, Austria) and mounted on formvar coated slot grids (Agar Scientific, Stansted, Essex, United Kingdom). They were contrasted with 0.5% aqueous uranyl acetate (Science Services, München, Germany) for 20 min and with 2% Reynold's lead citrate for 6 min before imaging at 20-30 kV with a Quanta FEG 250 scanning electron microscope (FEI Company, Hillsboro,

OR, USA) equipped with a STEM detector using the xT microscope control software v. 6.2.6.3123.

### Supplementary Figures

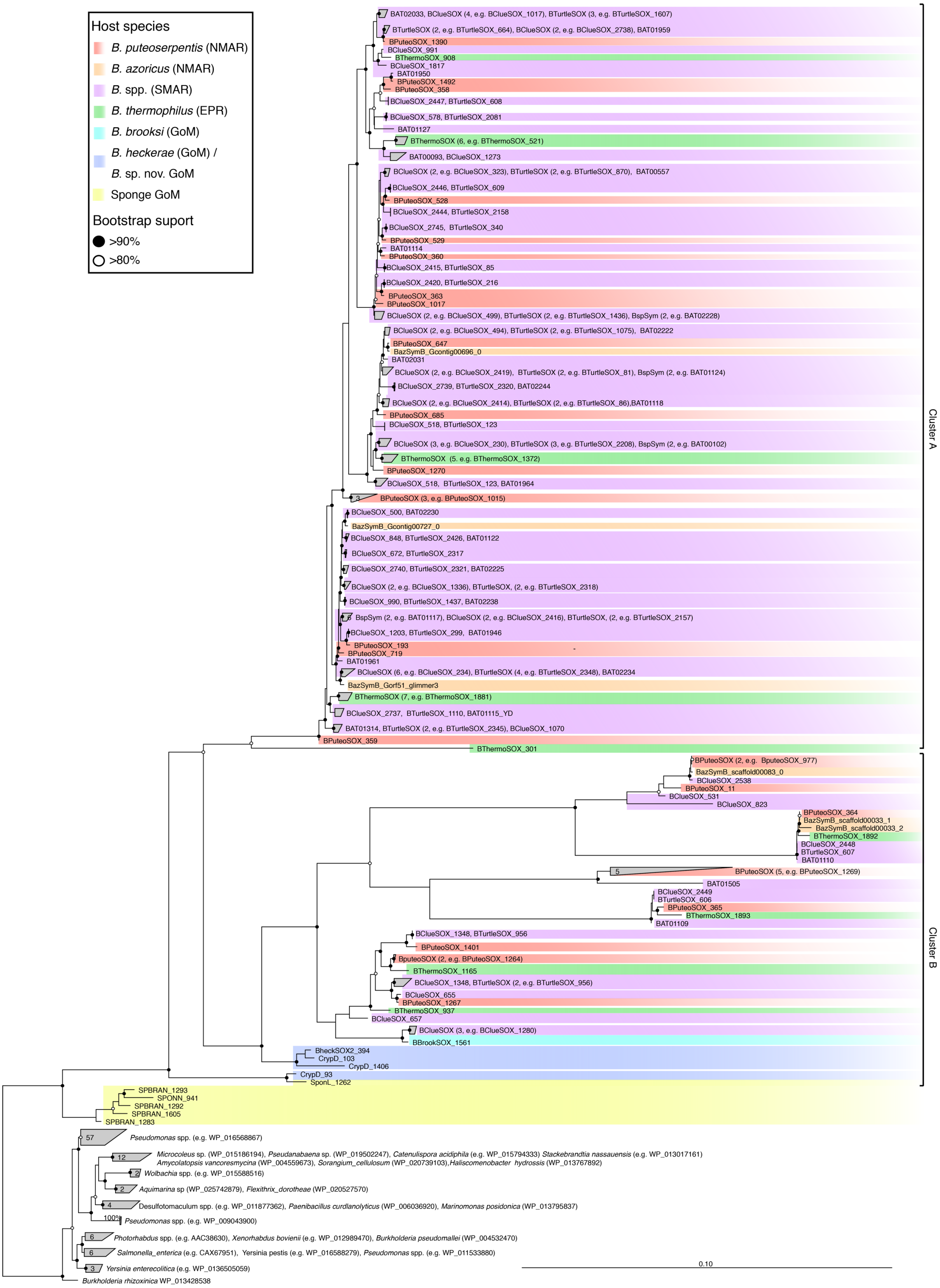

**Fig. S1.** Phylogeny of the YD repeat genes from SOX symbionts. The tree was calculated with the YD sequences of free-living and host-associated bacteria that had similar YD repeats to the SOX symbiont. Branches were collapsed into clusters if all sequences in the cluster came from symbionts from the same host species. The number of sequences in each cluster are indicated in parentheses. Locus tags of the YD genes from the SOX symbionts of *Bathymodiolus* sp., Clueless (BClueSOX), *B*. sp., Wide Awake (BTurtleSOX), *B*. sp. Lilliput (BAT), *B. puteoserpentis* (BPuteoSOX), *B. thermophilus* (BThermoSOX), *B. azoricus* (BazSymB), encrusting sponge from Mictlan, GoM (SponL), encrusting sponge, Campeche, GoM (SPONN), branching sponge, Chapopote, GoM (SPBRAN) are shown at the nodes.

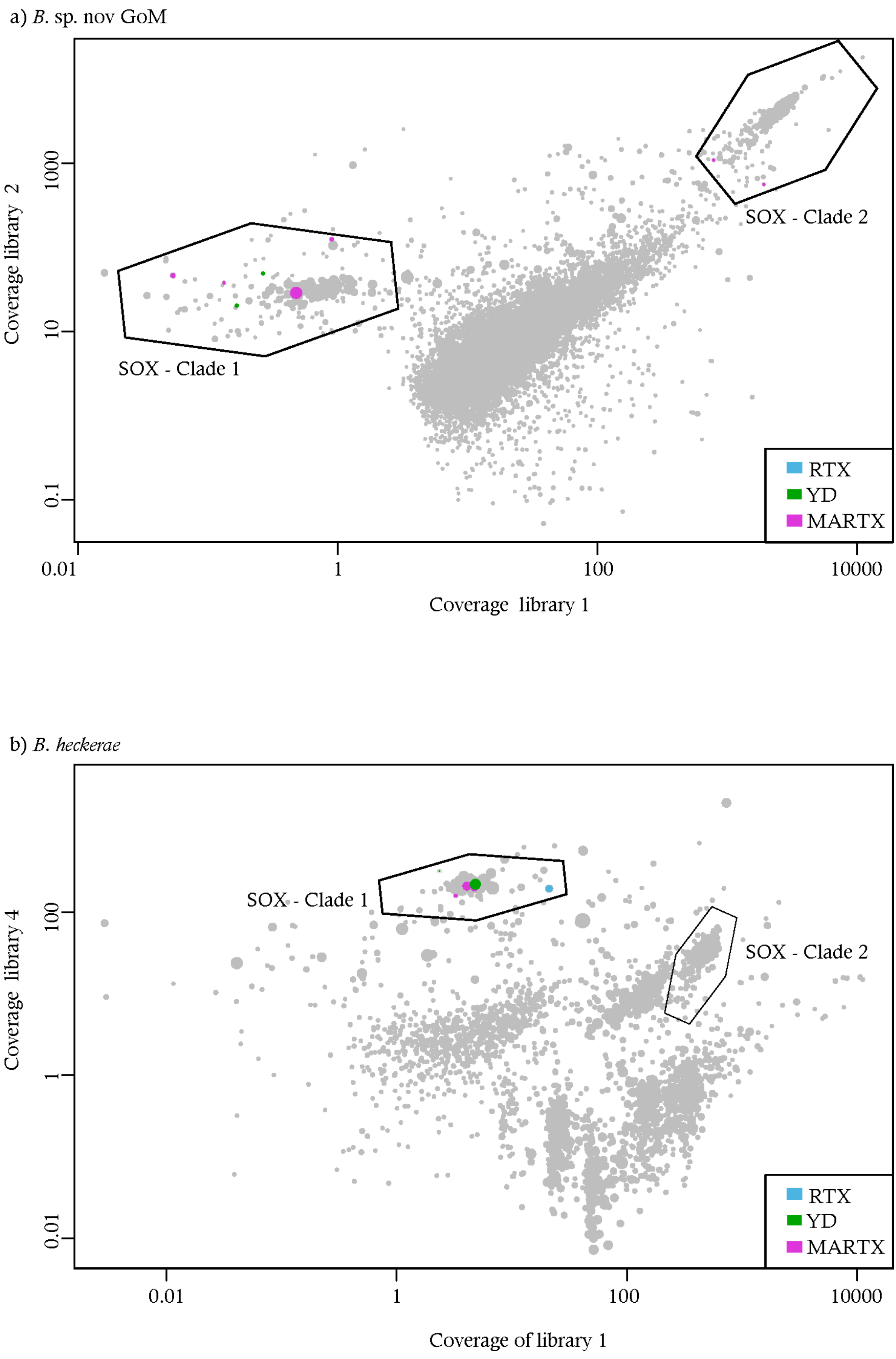
a) B. sp. nov GoM
Coverage library 2
1000
10
0.1
0.01
1
100
10000
Coverage library 1
SOX - Clade 1
SOX - Clade 2
RTX
YD
MARTX
b) B. heckerae
Coverage library 4
100
1
0.01
0.01
1
100
10000
Coverage of library 1
SOX - Clade 1
SOX - Clade 2
RTX
YD
MARTX

**Fig. S2** Differential sequencing coverage of the metagenome assemblies from *B*. sp. nov GoM (a) and *B. heckerae* (b). Based on the sequencing coverage, the YD genes and RTX present in the metagenome assemblies belong to the SOX symbionts that fall in Clade 1 (Fig. 1). Each dot represents one contig, and the size of the dots are scaled to the scaffold length. Only scaffolds longer than either 1000 bp for *B*. sp. nov GoM, or 2000 bp for *B. heckerae* are shown. A higher minimum scaffold length was used for *B. heckerae* to improve the differential coverage visualization as this mussel hosts more symbionts

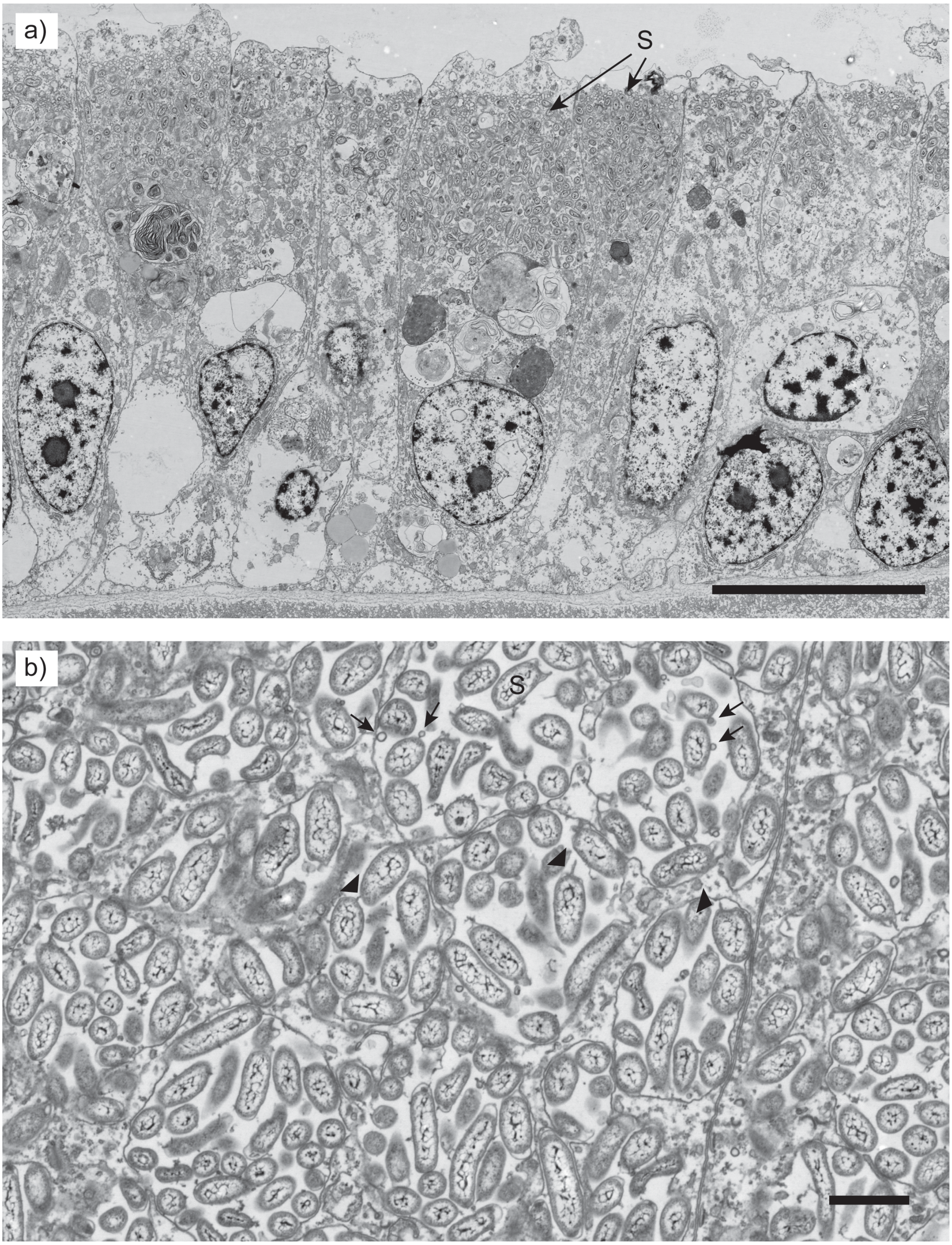


**Fig. S3.** Transmission electron micrographs of cross-sections of *B*. sp. nov. GoM gill cells. a) SOX symbionts (S) are located inside host vacuoles in the apical region of the bacteriocytes. Scale = 10 µm. b) Higher-magnification image of vacuoles containing SOX symbionts. Some of the symbionts appear to produce outer membrane vesicles (forming stage = arrowheads; complete stage = arrows). Scale = 1µm.

### Supplementary Tables

**Supplementary Table 1.** Samples sequenced and/or assembled in this study. Symbiont genomes and sequencing reads were submitted to the European Nucleotide Archive (ENA) repository under project accession numbers PRJEB17996 and PRJEB28154 for mussels and PRJEB20503 for sponges. MAR = Mid-Atlantic Ridge; EPR = East Pacific Rise; GoM = Gulf of Mexico.

| Species | Cruise | Collection date | Site | Latitude | Longitude | Depth (m) | Max. no. of sequencing reads | Acc. Number of symbiont genomes | # | Host CO1 |
|---|---|---|---|---|---|---|---|---|---|---|
| *B. thermophilus* | AT26-23 Dive 4763-2014 | 08.11.2014 | Crab-Spa, EPR | 9°50.377'N | 104°17.533'W | 2512 | 16.6[+,1,3] | ERS1452907 / SAMEA4553728 | 3 | KU597627 |
| *B. puteoserpentis* | M64-2 | 17.05.05 | Logatchev, MAR | 14° 45.3'N | 44° 59.2662'W | 3009 | 217.6[*,1,3]<br>120.2[*,1,4] | ERS1452910 / SAMEA4553731 | 1 | KU597632 |
| *B.* sp. 5° South, Clueless | M78-2 302 ROV15 | 22.04.2009 | 5° South, Clueless, MAR | 4° 48.19599'S | 12° 22.308024'W | 2971.5 | 15[+,2,5] | ERS1452938 SAMEA4553759 | 3 | LT674164 |
| *B.* sp. 5° South, Wide Awake | ATA 57 ROV 7/2 | 21.01.2008 | 5° South, Wide Awake, MAR | 4°48.558'S | 12°22.463'W | 2989 | 13.4[+,1,6] | ERS1453684 SAMEA4554505 | 1 | LT674165 |
| *B. heckerae* | M114-2 | 14.03.2015 | Chapopote, GoM | 21° 54.003'N | 93° 26.124'W | 2925 | 30.6[+,2,3] | **Clade 2 SOX symbiont:** ERS1453771 / SAMEA4554592 | 3 | LT674166 |
| *B. heckerae* | M67/2 | 13.04.2006 | Chapopote, GoM | 21° 53.98'N | 93° 26.12'W | 2923 | 23.1[+,2,5] | **Clade 1 SOX symbiont:** ERS1673412 / SAMEA103984244 | 1 | LT841270 |
| *B. brooksi* | M114-2 | 14.03.2015 | Chapopote, GoM | 21° 54.003'N | 93° 26.124'W | 2925 | 46.6[+,2,5] | ERS1453772 SAMEA4554593 | 3 | LT674167 |
| *B.* sp. *nov.* GoM | NA043-H1337 | 28.06.2014 | DC673, GoM | 28°18.5595'N | 87°18.6512'W | 2604 | 35.5[+,2,5] | **Clade 2 SOX symbiont:** ERS1673800 / SAMEA103984632<br>**Clade 1 SOX symbiont:** ERS1673801 / SAMEA103984633 | 2 | LT841271 |
| Encrusting sponge, Mictlan, GoM (ESM_GoM)[%] | M114-2 | 11.03.2015 | Mictlan, GoM | 22°1'N | 93°14'W | 3106 | 27.7[%,5] | ERS1673799/ SAMEA103984631 | 1 | KU659136-6 |
| Encrusting sponge, Chapopote, GoM (ESC_GoM)[%] | M114-2 | 20.03.2015 | Chapopote, GoM | 21° 54.003'N | 93° 26.124'W | 2925 | 42.6[%,5] | ERS1673797/ SAMEA103984629 | 1 | KU659136-7 |
| Branching sponge, Chapopote, GoM (BSC_GoM)[%] | M114-2 | 20.03.2015 | Chapopote, GoM | 21° 54.003'N | 93° 26.124'W | 2925 | 42.6[%,] 5 | ERS1673798/ SAMEA103984630 | 1 | KU659136-8 |

Sequencing center = [+]Max Planck Genome Centre (Germany); [&]University of Bielefeld (Germany); [*]Genoscope (France).

[%]Sequencing was done by Rubin-Blum *et al*., (2017).

#Number of individuals sequenced (independent libraries).

[1]DNA extraction as described by Zhou *et al*. (1996); [2]DNA extraction with DNAsy Blood and Tissue kit; [3]Sequenced as 2x 100 bp paired-end reads on an Illumina HiSeq 2000 instrument; [4]Sequenced as

2x 100 bp mate-paired reads of a 5.6 kb library on an Illumina HiSeq 2000 instrument; [5]Sequenced as 2x 150 bp paired-end reads on an Illumina HiSeq 2500 instrument; [6]Sequenced as 2x 250 bp paired-end reads on an Illumina MiSeq instrument.

**Supplementary Table 2.** Description of the domains found in MARTX genes from Clade 2 mussel SOX symbionts and the most similar MARTX sequences in other mussel SOX symbionts from Clade 1 that are shown in Fig. 3.

| Accession | Short name | Definition from InterPro (www.ebi.ac.uk/interpro) |
|---|---|---|
| pfam05860 | Haemagg_act | Haemagglutination activity domain; this domain is suggested to be a carbohydrate- dependent haemagglutination activity site. It is found in a range of haemagglutinins and haemolysins |
| COG2911 | TamB | Autotransporter translocation and assembly factor *tamB* (intracellular trafficking, secretion, and vesicular transport) |
| pfam13332 | Fil_haemagg_2 | Haemagluttinin repeat |
| pfam04830 | DUF637 | Possible hemagglutinin; This family represents a conserved region found in a bacterial protein which may be a hemagglutinin or hemolysin |
| pfam09000 | Cytotoxic | Cytotoxic; The cytotoxic domain confers cytotoxic activity to proteins, enabling the formation of nucleolytic breaks in 16S ribosomal RNA |
| cd00519 | Lipase_3 | Lipase (class 3). Lipases are esterases that can hydrolyze long-chain acyl-triglycerides into di- and monoglycerides, glycerol, and free fatty acids at a water/lipid interface |
| COG2931 | COG2931 | Ca2+-binding protein, RTX toxin-related |
| pfam08548 | Peptidase_M10_C | Peptidase M10 serralysin C terminal |
| cd00741 | Lipase | Lipase. Lipases are esterases that can hydrolyze long-chain acyl-triglycerides into di- and monoglycerides, glycerol, and free fatty acids at a water/lipid interface |
| PRK13914 | PRK13914 | Invasion associated secreted endopeptidase |
| pfam11713 | Peptidase_C80 | Peptidase C80 family |
| pfam03382 | DUF285 | DUF285. This region appears distantly related to leucine rich repeats |
| pfam13205 | Big_5 | Bacterial Ig-like domain |
| pfam13753 | SWM_repeat | Putative flagellar system-associated repeat |
| cd06582 | TM_PBP1_LivH_like | Transmembrane subunit (TM) of *Escherichia coli* LivH. LivH is one of two TMs of the *E. coli* LIV-1/LS transporter, an ATP-Binding Cassette (ABC) transporter involved in the uptake of branched-chain amino acids (AAs) |
| pfam12319 | TryThrA_C | Tryptophan-Threonine-rich *Plasmodium* antigen C terminal. This protein is common to eukaryotes. This family is the C terminal of a surface antigen of malarial *Plasmodium* species. |
|  | 34 | Long tail fiber, proximal subunit |
| pfam13517 | VCBS | Repeat domain in *Vibrio*, *Colwellia*, *Bradyrhizobium* and *Shewanella*. The large protein size and repeat copy numbers, species distribution, and suggested activities of several member proteins suggests a role for this domain in adhesion (TIGR) |
| PTZ00474 | PTZ00474 | tryptophan/threonine-rich antigen superfamily |
|  | T1SS_VCA0849 | Type I secretion C-terminal target domain (VC_A0849 subclass) |
| COG3210 | FhaB | Large exoprotein involved in heme utilization or adhesion (intracellular trafficking, secretion, and vesicular transport) |
| pfam11068 | YlqD | YlqD protein. This family may act as molecular chaperones. |
| cd11304 | Cadherin_repeat | Cadherins are glycoproteins involved in Ca2+-mediated cell-cell adhesion. They play numerous roles in cell fate, signaling, proliferation, differentiation, and migration |
| pfam03382 | DUF285 | DUF285. This region appears distantly related to leucine rich repeats |

**Supplementary Table 3.** Secretion systems present in the SOX symbionts and their close-relatives. VirB5 and VirB11 were manually curated. If a secretion system class was described as having more genes than those detected in the genomes analyzed in this study, the maximum number of genes detected out of the total mandatory genes is shown. Mandatory genes are defined as always present in the system, accessory as some times present, and forbidden as never present (Abby *et al*., 2016). Type IV pilus (T4P); Tight adherence (Tad).

| Class | Transporter | ***Bathymodiolus* symbionts** | | | | | | | | | | | | **Sponge symbionts** | | | **Vesicomyid symbionts** | | **Free-living** | | |
|---|---|---|---|---|---|---|---|---|---|---|---|---|---|---|---|---|---|---|---|---|---|
| | | ***B. puteoserpentis* sym** | ***B. azoricus* sym[2]** | ***B.* sp. Lilliput sym[1]** | ***B.* sp. Clueles sym** | ***B.* sp. Wide Awake sym** | ***B. brooksi* sym** | ***B. thermophilus* sym** | ***B.* sp. nov. GoM** | ***B. heckerae* sym** | ***B. septemdierum* sym[3]** | ***B.* sp. nov GoM sym** | ***B. heckerae* sym** | **ESM_GoM** | **ESC_GoM** | **BSC_GoM** | ***Ca.* R. magnifica[4]** | ***Ca.* V. okutanii[5]** | ***Ca.* T. singularis[6]** | ***Ca.* T. autotrophicus[7]** | **SUP05[8]** |
| | | **Clade 1** | | | | | | | | | | **Clade 2** | | **Clade 1** | | | **-** | | **Clade2** | | |
| T1SS, mandatory | T1SS_abc | 6 | 5 | 5 | 6 | 5 | 5 | 6 | 5 | 5 | 3 | 3 | 3 | 5 | 5 | 3 | 3 | 2 | 3 | 3 | 3 |
| T1SS, mandatory | T1SS_mfp | 2 | 1 | 2 | 2 | 2 | 2 | 3 | 2 | 2 | 1 | | | 1 | 1 | | 1 | | | | |
| T1SS, mandatory | T1SS_omf | 2 | 2 | 2 | 2 | 2 | 2 | 2 | 3 | 3 | 2 | 1 | 1 | 2 | 3 | 2 | 2 | | | 2 | 1 |
| T2SS, 8 of 9 mandatory | T2SS_gspD | 1 | 1 | | | | 1 | 1 | | | | | | | | | | | | | |
| T2SS, 8 of 9 mandatory | T2SS_gspE | 1 | 1 | 1 | 1 | 1 | 1 | 1 | 1 | 1 | | | | 1 | 1 | 1 | | | | | |
| T2SS, 8 of 9 mandatory | T2SS_gspF | 1 | 1 | 1 | 1 | 1 | 1 | 1 | 1 | 1 | | | | 1 | 1 | 1 | | | | | |
| T2SS, 8 of 9 mandatory | T2SS_gspG | | 2 | 2 | 1 | 1 | 1 | 1 | 1 | 1 | | | | 1 | 1 | 1 | | | 1 | | |
| T2SS, 8 of 9 mandatory | T2SS_gspH | | | | | | 1 | 1 | 1 | 1 | | | | 1 | 1 | | | | | | |
| T2SS, 8 of 9 mandatory | T2SS_gspI | 1 | 1 | 1 | 1 | 1 | 1 | 1 | 1 | 1 | | | | 1 | 1 | 1 | | | | 1 | 1 |
| T2SS, 8 of 9 mandatory | T2SS_gspJ | 1 | 1 | 1 | 1 | 1 | 1 | 1 | 1 | 1 | | | | 1 | 1 | 1 | | | | 2 | 1 |
| T2SS, 8 of 9 mandatory | T2SS_gspK | 1 | | 1 | 1 | 1 | | 1 | 1 | 1 | | | | 1 | 1 | 1 | | | | | |
| T2SS, accessory | T2SS_gspC | 1 | 1 | 1 | 1 | 1 | 1 | 1 | 1 | 1 | | | | 1 | 1 | 1 | | | | | |
| T2SS, accessory | T2SS_gspN | | | | | | 1 | 1 | 1 | 1 | | | | | | 1 | | | | | |
| T3SS, 1 of 8 mandatory | T3SS_sctN | 2 | 2 | 2 | 2 | 2 | 2 | 2 | 1 | 1 | 1 | 2 | 2 | 1 | 1 | 1 | 2 | 2 | 2 | 2 | 1 |
| T4P, accessory | T4P_pilD | 1 | 1 | 1 | 1 | 1 | 1 | 1 | 1 | 1 | | | | | | 1 | | | 1 | 1 | 1 |
| T4P, 7 of 10 mandatory | T4P_pilAE | | | | 1 | 1 | | | | | | | | | | | | | 4 | | 3 |
| T4P, 7 of 10 mandatory | T4P_pilB | | 1 | 1 | 1 | 1 | | 1 | | | | | | | | | | | 1 | 1 | 1 |
| T4P, 7 of 10 mandatory | T4P_pilC | | | | | | | | | | | | | | | | | | 1 | 1 | 1 |
| T4P, 7 of 10 mandatory | T4P_pilO | | | | | | | | | | | | | | | | | | 1 | 1 | |
| T4P, 7 of 10 mandatory | T4P_pilQ | | 1 | 2 | 2 | 2 | | 1 | 1 | 1 | 1 | | | 1 | 1 | 1 | | | 1 | 1 | |
| T4P, 7 of 10 mandatory | T4P_pilT_pilU | | | 1 | 1 | 1 | | | | | | | | | | | | | 1 | 1 | 1 |
| T4P, 7 of 10 mandatory; **Forbidden in T2SS** | T4P_pilM | 1 | 1 | 1 | 1 | 1 | 1 | 1 | 1 | 1 | | 1 | 1 | | | 1 | 1 | | 1 | 1 | 1 |
| **T4SS_typeI, F, G and B, mandatory** | T4SS_MOBB | | 1 | 2 | | 1 | | | | | | 1 | 2 | | | | | | | | |
| **T4SS_typeI, F, G and B, mandatory** | T4SS_t4cp1 | | 3 | 2 | | 1 | | | | | | 1 | 2 | | | | | | | | |
| **T4SS_typeI, F, G and B, mandatory** | T4SS_virb4 | | 1 | 1 | | 1 | | | | | | 1 | 2 | | | | | | | | |
| T4SS_typeB, accessory | T4SS_B_traP | | | | | | | | | | | 1 | 1 | | | | | | | | |
| T4SS_typeF, accessory | T4SS_F_traB | | | | | | | | | | | 1 | 1 | | | | | | | | |
| T4SS_typeF, accessory | T4SS_F_traE | | 1 | | | | | | | | | 1 | 1 | | | | | | | | |
| T4SS_typeF, accessory | T4SS_F_traF | | 1 | | | | | | | | | 1 | 1 | | | | | | | | |
| T4SS_typeF, accessory | T4SS_F_traG | | | | | | | | | | | | | | | | | | | | |
| T4SS_typeF, accessory | T4SS_F_traH | | 1 | | | | | | | | | 1 | 1 | | | | | | | | |
| T4SS_typeF, accessory | T4SS_F_traK | | | | | | | | | | | 1 | 1 | | | | | | | | |
| T4SS_typeF, accessory | T4SS_F_traL | | 1 | | | | | | | | | | | | | | | | | | |
| T4SS_typeF, accessory | T4SS_F_traU | | 1 | | | | | | | | | 1 | 1 | | | | | | | | |
| T4SS_typeF, accessory | T4SS_F_traV | | 1 | | | | | | | | | 1 | 1 | | | | | | | | |
| T4SS_typeF, accessory | T4SS_F_traW | | 1 | | | | | | | | | 1 | 1 | | | | | | | | |
| T4SS_typeF, accessory | T4SS_G_tfc7 | | 1 | 1 | | 1 | | | | | | | | | | | | | | | |
| T4SS_typeF, accessory | T4SS_F_trbC | | 1 | | | | | | | | | 1 | 1 | | | | | | | | |
| T4SS_typeG, accessory | T4SS_G_tfc12 | | 1 | 1 | | 1 | | | | | | | | | | | | | | | |
| T4SS_typeG, accessory | T4SS_G_tfc13 | | | 1 | | 1 | | | | | | | | | | | | | | | |
| T4SS_typeG, accessory | T4SS_G_tfc14 | | 1 | 1 | | 1 | | | | | | | | | | | | | | | |
| T4SS_typeG, accessory | T4SS_G_tfc15 | | 1 | | | | | | | | | | | | | | | | | | |
| T4SS_typeG, accessory | T4SS_G_tfc19 | | | | | 1 | | | | | | | | | | | | | | | |
| T4SS_typeG, accessory | T4SS_G_tfc2 | | | 1 | 1 | 1 | | 1 | | | | | | | | | | | | | |
| T4SS_typeG, accessory | T4SS_G_tfc22 | | | 1 | | 1 | | | | | | | | | | | | | | | |
| T4SS_typeG, accessory | T4SS_G_tfc23 | | 1 | 1 | | 1 | | | | | | | | | | | | | | | |
| T4SS_typeG, accessory | T4SS_G_tfc24 | | 1 | 1 | | 1 | | | | | | | | | | | | | | | |
| T4SS_typeG, accessory | T4SS_G_tfc3 | | 1 | | | 1 | | | | | | | | | | | | | | | |
| T4SS_typeG, accessory | T4SS_G_tfc5 | | 1 | 1 | | 1 | | | | | | | | | | | | | | | |
| T4SS_typeG, accessory | T4SS_G_tfc8 | | | 1 | | 1 | | | | | | | | | | | | | | | |
| T4SS_typeI, accessory | T4SS_I_traM | | | | | | | | | | | | 1 | | | | | | | | |

| Class | Transporter | ***Bathymodiolus* symbionts** | | | | | | | | | | | | **Sponge symbionts** | | | **Vesicomyid symbionts** | | **Free-living** | | |
|---|---|---|---|---|---|---|---|---|---|---|---|---|---|---|---|---|---|---|---|---|---|
| | | ***B. puteoserpentis* sym** | ***B. azoricus* sym[2]** | ***B.* sp. Lilliput sym[1]** | ***B.* sp. Clueles sym** | ***B.* sp. Wide Awake sym** | ***B. brooksi* sym** | ***B. thermophilus* sym** | ***B.* sp. nov. GoM** | ***B. heckerae* sym** | ***B. septemdierum* sym[3]** | ***B.* sp. nov GoM sym** | ***B. heckerae* sym** | **ESM_GoM** | **ESC_GoM** | **BSC_GoM** | ***Ca.* R. magnifica[4]** | ***Ca.* V. okutanii[5]** | ***Ca.* T. singularis[6]** | ***Ca.* T. autotrophicus[7]** | **SUP05[8]** |
| | | **Clade 1** | | | | | | | | | | **Clade 2** | | **Clade 1** | | | **-** | | **Clade2** | | |
| T4SS_typeI, accessory | T4SS_I_traP | | | 1 | | 1 | | | | | | | | | | | | | | | |
| T4SS_typeT, accessory | T4SS_T_virB10 | | | | | | | | | | | | 1 | | | | | | | | |
| T4SS_typeT, accessory | T4SS_T_virB11 | | | | | | | | | | | | 1 | | | | | | | | |
| T4SS_typeT, accessory | T4SS_T_virB2 | | | | | | | | | | | | 1 | | | | | | | | |
| T4SS_typeT, accessory | T4SS_T_virB3 | | | | | | | | | | | | 1 | | | | | | | | |
| T4SS_typeT, accessory | T4SS_T_virB5 | | | | | | | | | | | | 1 | | | | | | | | |
| T4SS_typeT, accessory | T4SS_T_virB6 | | | | | | | | | | | | 1 | | | | | | | | |
| T4SS_typeT, accessory | T4SS_T_virB8 | | | 1 | | | | | | | | | 1 | | | | | | | | |
| T4SS_typeT, accessory | T4SS_T_virB9 | | | | | | | | | | | | 1 | | | | | | | | |
| T5SS, mandatory | T5aSS_PF03797 | 2 | 4 | 1 | 5 | 4 | | 1 | 1 | | | | | 1 | 1 | 1 | | | 1 | | 1 |
| T5SS, mandatory | T5bSS_translocator, also an RTX activator | 2 | | 2 | 2 | 2 | | | | | | | | | | | | | | | |
| T5SS, mandatory | T5cSS_PF03895 | | | | | | | | | | | | 1 | | | 1 | | | | | 1 |
| T6SSi, 1 of 10 mandatory | T6SSi_tssH | 1 | 1 | 1 | 1 | 1 | 1 | 1 | 1 | 1 | 1 | 1 | 1 | 1 | 1 | 1 | 1 | 1 | | 1 | 1 |
| Tad, 2 of 6 mandatory | Tad_tadA | | | | | | | | | | | | 1 | | | | | | | | |
| Tad, 2 of 6 mandatory; **forbidden in T4P** | Tad_tadZ | | | | | | | | | 1 | 1 | | | | | | | | | | |
| Flagellum, 1 of 8 mandatory | Flg_sctN_FLG | 1 | 1 | 1 | 1 | 1 | 1 | 1 | 2 | 2 | 2 | 1 | 1 | 2 | | 2 | 1 | 1 | 1 | 1 | 1 |

Accession numbers: [1]PRJNA65421 and IMG 2518645510; [2]PRJEB8263, [3]NZ_AP013042; [4]NC_008610; [5]NC_009465; [6]CP006911; [7]NZ_CP010552; [8]ACSG00000000

**Supplementary Table 4.** Genome statistics of SOX symbiont genomes from individuals sequenced from the same geographic sites as described in Supplementary Table 1. The number of TRGs in these genomes was similar to the representative genomes shown in Fig. 1.

| Host species | Cruise | Collection date | Site | Latitude | Longitude | Completeness | YD | RTX | MARTX | Genome sequencing coverage |
|---|---|---|---|---|---|---|---|---|---|---|
| *B. thermophilus** | AT26-23 Dive 4763-2014 | 08.11.14 | Crab-Spa, EPR | 9°50.377'N | 104°17.533'W | 97.19% | 27 | 1 | 15 | 779 |
| *B. thermophilus** | AT26-23 Dive 4763-2014 | 08.11.14 | Crab-Spa, EPR | 9°50.377'N | 104°17.533'W | 97.85% | 30 | 1 | 17 | 477 |
| *B*. sp. 5° South, Clueless* | M78-2 302, ROV15 | 22.04.09 | 5° South, Clueless, MAR | 4° 48.19599'S | 12° 22.308024'W | 98.51% | 65 | 2 | 33 | 398 |
| *B*. sp. 5° South, Clueless* | M78-2 302, ROV15 | 22.04.09 | 5° South, Clueless, MAR | 4° 48.19599'S | 12° 22.308024'W | 98.51% | 59 | 0 | 33 | 778 |
| *B. heckerae*** | M114-2 | 14.03.15 | Chapopote, GoM | 21° 54.003'N | 93° 26.124'W | 97.22% | 0 | 0 | 1 | 63 |
| *B. heckerae*** | M114-2 | 14.03.15 | Chapopote, GoM | 21° 54.003'N | 93° 26.124'W | 97.22% | 0 | 0 | 1 | 106 |
| *B. brooksi** | M114-2 | 14.03.15 | Chapopote, GoM | 21° 54.003'N | 93° 26.124'W | 97.19% | 1 | 0 | 3 | 17 |
| *B. brooksi** | M114-2 | 14.03.15 | Chapopote, GoM | 21° 54.003'N | 93° 26.124'W | 97.19% | 1 | 0 | 3 | 12 |
| *B*. sp. *nov*. GoM** | NA043-H1337 | 28.06.14 | DC673, GoM | 28°18.5595'N | 87°18.6512'W | 98.68% | 0 | 0 | 3 | 234 |
| *B*. sp. *nov*. GoM* | NA043-H1337 | 28.06.14 | DC673, GoM | 28°18.5595'N | 87°18.6512'W | 0% | NA | NA | NA | 3 |

*Clade 1 SOX symbiont

**Clade 2 SOX symbiont

**Supplementary Table 5.** Domain scores for classifying genes as MARTX, RTX or YD. The protein was only classified as a TRG when the total domain score was higher than or equal to one. Some domains present in TRGs can disrupt the normal functioning of the cell by having similar domains to proteins involved in core metabolic functions. Thus, we used the term 'in combination' for domains that were also present in proteins with putative functions in core cellular metabolism.

| Accession | Short name | Definition (from InterPro database) | MARTX | RTX | YD | RTX activator | Class |
|---|---|---|---|---|---|---|---|
| cl00296 | Peptidase_C39_like superfamily | Peptidase family C39 mostly contains bacteriocin-processing endopeptidases from bacteria | 0 | 0.5 | 0 | 0 | RTX? |
| PHA02584 | 34 | Long tail fiber, proximal subunit | 0 | 1 | 0 | 0 | RTX (in combination) |
| cl21494 | Esterase_lipase superfamily | Esterases and lipases (includes fungal lipases, cholinesterases, etc.) | 0 | 0.5 | 0 | 0 | RTX (in combination) |
| pfam13529 | Peptidase_C39_2 | Peptidase_C39 like family | 0 | 1 | 0 | 0 | RTX (in combination) |
| COG5153 | CVT17 | Putative lipase essential for disintegration of autophagic bodies inside the vacuole | 0 | 2 | 0 | 0 | RTX |
| cl16721 | DUF4329 superfamily | Domain of unknown function (DUF4329) | 0 | 0 | 0.1 | 0 | YD (in combination) |
| cl17169 | RRM_SF superfamily | RNA recognition motif (RRM) superfamily | 0 | 0 | 0.1 | 0 | YD (in combination) |
| cl15554 | SpvB superfamily | Salmonella virulence plasmid 65kda B protein | 0 | 0 | 0.1 | 0 | YD (in combination) |
| pfam13517 | VCBS | Repeat domain in *Vibrio*, *Colwellia*, *Bradyrhizobium* and *Shewanella*. The large protein size and repeat copy numbers, species distribution, and suggested activities of several member proteins suggests a role for this domain in adhesion (TIGR) | 1 | 0 | 1 | 0 | YD (in combination) |
| PRK15244 | PRK15244 | Virulence protein spvb | 0 | 0 | 2 | 0 | YD |
| TIGR03696 | Rhs_assc_core | RHS repeat-associated core domain. This class includes secreted bacterial insecticidal toxins and intercellular signaling proteins such as teneurins in animals | 0 | 0 | 2 | 0 | YD |
| cl14012 | Rhs_assc_core superfamily | RHS repeat-associated core domain | 0 | 0 | 2 | 0 | YD |
| pfam05593 | RHS_repeat | Rhs repeat | 0 | 0 | 2 | 0 | YD |
| cl11982 | RHS_repeat superfamily | Rhs repeat | 0 | 0 | 2 | 0 | YD |
| COG3209 | RhsA | Uncharacterized conserved protein rhas | 0 | 0 | 2 | 0 | YD |
| pfam03534 | SpvB | Salmonella virulence plasmid 65kda B protein | 0 | 0 | 2 | 0 | YD |

| Accession | Short name | Definition (from InterPro database) | MARTX | RTX | YD | RTX activator | Class |
|---|---|---|---|---|---|---|---|
| pfam12255 | TcdB_toxin_midC | Insecticide toxin tcdb middle/C-terminal region | 0 | 0 | 2 | 0 | YD |
| cl13662 | TcdB_toxin_midC superfamily | Insecticide toxin tcdb middle/C-terminal region | 0 | 0 | 2 | 0 | YD |
| pfam12256 | TcdB_toxin_midN | Insecticide toxin tcdb middle/N-terminal region | 0 | 0 | 2 | 0 | YD |
| cl13663 | TcdB_toxin_midN superfamily | Insecticide toxin tcdb middle/N-terminal region | 0 | 0 | 2 | 0 | YD |
| cl21563 | VCBS superfamily | Repeat domain in *Vibrio*, *Colwellia*, Bradyrhizobium and *Shewanella*. The large protein size and repeat copy numbers, species distribution, and suggested activities of several member proteins suggests a role for this domain in adhesion (TIGR) | 2 | 0 | 2 | 0 | YD |
| pfam03538 | VRP1 | *Salmonella* virulence plasmid 28.1kda A protein | 0 | 0 | 2 | 0 | YD |
| cl21676 | VRP1 superfamily | *Salmonella* virulence plasmid 28.1kda A protein | 0 | 0 | 2 | 0 | YD |
| TIGR01643 | YD_repeat_2x | YD repeat (two copies) | 0 | 0 | 2 | 0 | YD |
| cl00125 | RHOD superfamily | Rhodanese homology domain (rhod) | -10 | -10 | -10 | 0 | Rhodonase |
| TIGR02172 | Fb_sc_TIGR02172 | *Fibrobacter succinogenes* paralogous family TIGR02172 | 0.01 | 0.01 | 0 | 0 | MARTX/RTX, not specific |
| pfam11713 | Peptidase_C80 | Peptidase C80 family | 1.01 | 0.5 | 0 | 0 | MARTX/RTX, more common in MARTX |
| cl21453 | PKc_like superfamily | Protein kinases, catalytic domain | 0.05 | 0.05 | 0 | 0 | MARTX/RTX (in combination) |
| pfam03160 | Calx-beta | Calx-beta domain | 1 | 1 | 0 | 0 | MARTX/RTX |
| COG2931 | COG2931 | Ca2+-binding protein, RTX toxin-related | 2 | 2 | 0 | 0 | MARTX/RTX |
| pfam05557 | MAD | Mitotic checkpoint protein | 1 | 1 | 0 | 0 | MARTX/RTX |
| cl13207 | Peptidase_C80 superfamily | Peptidase C80 family | 1 | 1 | 0 | 0 | MARTX/RTX |
| pfam08548 | Peptidase_M10_C | Peptidase M10 serralysin C terminal | 0.6 | 0.5 | 0 | 0 | MARTX/RTX |
| pfam05701 | WEMBL | Regulates the movement of cp-actin filaments | 1 | 0 | 1 | 0 | MARTX/RHS |
| COG2831 | FhaC | Hemolysin activation/secretion | 0.01 | 0 | 0 | 1 | MARTX activator |
| pfam08479 | POTRA_2 | POTRA domain, shlb-type. Shlb is important in the secretion and activation of the haemolysin shla | 0.01 | 0.01 | 0 | 1 | MARTX activator |
| PHA02562 | 46 | Endonuclease subunit | 0.01 | 0 | 0 | 0 | MARTX (In combination) |
| TIGR01612 | 235kDa-fam | Reticulocyte binding/rhoptry protein | 0.01 | 0 | 0 | 0 | MARTX (In combination) |

| Accession | Short name | Definition (from InterPro database) | MARTX | RTX | YD | RTX activator | Class |
|---|---|---|---|---|---|---|---|
| pfam13166 | AAA_13 | AAA domain; this family of domains contains a P-loop motif that is characteristic of the AAA superfamily. Many of the proteins in this family are conjugative transfer proteins | 0.01 | 0 | 0 | 0 | MARTX (In combination) |
| pfam13514 | AAA_27 | AAA domain; this domain is found in a number of double-strand DNA break proteins | 0.01 | 0 | 0 | 0 | MARTX (In combination) |
| TIGR01901 | adhes_NPXG | Filamentous hemagglutinin family N-terminal domain. Members of this family have been characterized as adhesins, filamentous haemagglutinins, heme/hemopexin-binding protein, etc. | 0.01 | 0 | 0 | 0 | MARTX (In combination) |
| pfam04111 | APG6 | Autophagy protein Apg6; in yeast, 15 Apg proteins coordinate the formation of autophagosomes | 0.01 | 0 | 0 | 0 | MARTX (In combination) |
| pfam13754 | Big_3_4 | Bacterial Ig-like domain (group 3) | 0.01 | 0 | 0 | 0 | MARTX (in combination) |
| cl16375 | Big_3_4 superfamily | Bacterial Ig-like domain (group 3) | 0.01 | 0 | 0 | 0 | MARTX (in combination) |
| pfam13205 | Big_5 | Bacterial Ig-like domain | 0.01 | 0 | 0 | 0 | MARTX (in combination) |
| pfam05262 | Borrelia_P83 | *Borrelia* P83/100 protein | 0.01 | 0 | 0 | 0 | MARTX (In combination) |
| cd11304 | Cadherin_repeat | Cadherins are glycoproteins involved in Ca2+-mediated cell-cell adhesion. They play numerous roles in cell fate, signaling, proliferation, differentiation, and migration | 0.01 | 0 | 0 | 0 | MARTX (in combination) |
| pfam11600 | CAF-1_p150 | Chromatin assembly factor 1 complex p150 subunit, N-terminal | 0.01 | 0 | 0 | 0 | MARTX (In combination) |
| pfam07888 | CALCOCO1 | Calcium binding and coiled-coil domain (CALCOCO1) like | 0.01 | 0 | 0 | 0 | MARTX (In combination) |
| pfam10174 | Cast | RIM-binding protein of the cytomatrix active zone; this is a family of proteins that form part of the CAZ (cytomatrix at the active zone) complex, which is involved in determining the site of synaptic vesicle fusion | 0.01 | 0 | 0 | 0 | MARTX (In combination) |
| TIGR03346 | chaperone_ClpB | ATP-dependent chaperone clpb. This molecular chaperone does not act as a protease, but rather serves to | 0.01 | 0 | 0 | 0 | MARTX (In combination) |

| Accession | Short name | Definition (from InterPro database) | MARTX | RTX | YD | RTX activator | Class |
|---|---|---|---|---|---|---|---|
| | | disaggregate misfolded and aggregated proteins | | | | | |
| COG4372 | COG4372 | Uncharacterized conserved protein, contains DUF3084 domain | 0.01 | 0 | 0 | 0 | MARTX (In combination) |
| COG4913 | COG4913 | Uncharacterized protein | 0.01 | 0 | 0 | 0 | MARTX (In combination) |
| COG5281 | COG5281 | Phage-related minor tail protein | 0.01 | 0 | 0 | 0 | MARTX (In combination) |
| pfam03344 | Daxx | Daxx family; the Daxx protein (also known as the Fas-binding protein) is thought to play a role in apoptosis | 0.01 | 0 | 0 | 0 | MARTX (In combination) |
| pfam09756 | DDRGK | DDRGK domain | 0.01 | 0 | 0 | 0 | MARTX (In combination) |
| pfam07263 | DMP1 | Dentin matrix protein 1 (DMP1); this family consists of several mammalian dentin matrix protein 1 (DMP1) sequences | 0.01 | 0 | 0 | 0 | MARTX (In combination) |
| pfam07423 | DUF1510 | Protein of unknown function (DUF1510) | 0.01 | 0 | 0 | 0 | MARTX (In combination) |
| cl22487 | DUF1510 superfamily | Protein of unknown function (DUF1510) | 0.01 | 0 | 0 | 0 | MARTX (In combination) |
| pfam12128 | DUF3584 | Protein of unknown function (DUF3584) | 0.01 | 0 | 0 | 0 | MARTX (In combination) |
| pfam04094 | DUF390 | Protein of unknown function (DUF390) | 0.01 | 0 | 0 | 0 | MARTX (In combination) |
| pfam05667 | DUF812 | Protein of unknown function (DUF812) | 0.01 | 0 | 0 | 0 | MARTX (In combination) |
| pfam05917 | DUF874 | *Helicobacter pylori* protein of unknown function (DUF874) | 0.01 | 0 | 0 | 0 | MARTX (In combination) |
| COG4942 | EnvC | Septal ring factor envc, activator of murein hydrolases amia and amib | 0.01 | 0 | 0 | 0 | MARTX (In combination) |
| COG4477 | EzrA | Septation ring formation regulator ezra | 0.01 | 0 | 0 | 0 | MARTX (In combination) |
| pfam13332 | Fil_haemagg_2 | Haemagluttinin repeat | 0.01 | 0 | 0 | 0 | MARTX (In combination) |
| cl16241 | Fil_haemagg_2 superfamily | Haemagluttinin repeat | 0.01 | 0 | 0 | 0 | MARTX (In combination) |
| pfam05860 | Haemagg_act | Haemagglutination activity domain | 0.01 | 0 | 0 | 0 | MARTX (In combination) |
| smart00912 | Haemagg_act | Haemagglutination activity | 0.01 | 0 | 0 | 0 | MARTX (In combination) |
| cl05436 | Haemagg_act superfamily | Haemagglutination activity domain | 0.01 | 0 | 0 | 0 | MARTX (In combination) |
| pfam07111 | HCR | Alpha helical coiled-coil rod protein (HCR). The function of HCR is | 0.01 | 0 | 0 | 0 | MARTX (In combination) |

| Accession | Short name | Definition (from InterPro database) | MARTX | RTX | YD | RTX activator | Class |
|---|---|---|---|---|---|---|---|
| | | unknown but it has been implicated in psoriasis in humans and is thought to affect keratinocyte proliferation. | | | | | |
| PRK11448 | hsdR | Type I restriction enzyme ecoki subunit R | 0.01 | 0 | 0 | 0 | MARTX (In combination) |
| smart00283 | MA | Methyl-accepting chemotaxis-like domains (chemotaxis sensory transducer) | 0.01 | 0 | 0 | 0 | MARTX (In combination) |
| pfam09726 | Macoilin | Transmembrane protein | 0.01 | 0 | 0 | 0 | MARTX (In combination) |
| pfam05672 | MAP7 | MAP7 (E-MAP-115) family; the organisation of microtubules varies with cell type and is presumably controlled by tissue-specific microtubule-associated proteins (maps). The 115-kda epithelial MAP (E-MAP-115/MAP7) has been identified as a microtubule-stabilizing protein predominantly expressed in cell lines of epithelial origin | 0.01 | 0 | 0 | 0 | MARTX (In combination) |
| COG5271 | MDN1 | Midasin, AAA ATPase with vwa domain, involved in ribosome | 0.01 | 0 | 0 | 0 | MARTX (In combination) |
| TIGR04523 | Mplasa_alph_rch | Helix-rich *Mycoplasma* protein | 0.01 | 0 | 0 | 0 | MARTX (In combination) |
| COG3264 | MscK | Small-conductance mechanosensitive channel | 0.01 | 0 | 0 | 0 | MARTX (In combination) |
| COG3096 | MukB | Chromosome condensin mukbef, ATPase and DNA-binding subunit mukb Uncharacterized protein involved in chromosome partitioning | 0.01 | 0 | 0 | 0 | MARTX (In combination) |
| PRK04863 | mukB | Cell division protein mukb | 0.01 | 0 | 0 | 0 | MARTX (In combination) |
| pfam05616 | Neisseria_TspB | *Neisseria meningitidis* tspb protein | 0.01 | 0 | 0 | 0 | MARTX (In combination) |
| pfam10168 | Nup88 | Nuclear pore component; Nup88 is overexpressed in tumor cells | 0.01 | 0 | 0 | 0 | MARTX (In combination) |
| PRK00247 | PRK00247 | Putative inner membrane protein translocase component yidc | 0.01 | 0 | 0 | 0 | MARTX (In combination) |
| PRK00409 | PRK00409 | Recombination and DNA strand exchange inhibitor protein | 0.01 | 0 | 0 | 0 | MARTX (In combination) |
| PRK02224 | PRK02224 | Chromosome segregation protein | 0.01 | 0 | 0 | 0 | MARTX (In combination) |
| PRK03918 | PRK03918 | Chromosome segregation protein | 0.01 | 0 | 0 | 0 | MARTX (In combination) |
| PRK05035 | PRK05035 | Electron transport complex protein rnfc | 0.01 | 0 | 0 | 0 | MARTX (In combination) |

| Accession | Short name | Definition (from InterPro database) | MARTX | RTX | YD | RTX activator | Class |
|---|---|---|---|---|---|---|---|
| PRK05771 | PRK05771 | V-type ATP synthase subunit I | 0.01 | 0 | 0 | 0 | MARTX (In combination) |
| PRK06975 | PRK06975 | Bifunctional uroporphyrinogen-III synthetase/uroporphyrin-III C-methyltransferase | 0.01 | 0 | 0 | 0 | MARTX (In combination) |
| PRK08581 | PRK08581 | N-acetylmuramoyl-L-alanine amidase | 0.01 | 0 | 0 | 0 | MARTX (In combination) |
| PRK09418 | PRK09418 | Bifunctional 2',3'-cyclic nucleotide 2'-phosphodiesterase/3'-nucleotidase precursor protein | 0.01 | 0 | 0 | 0 | MARTX (In combination) |
| PRK11281 | PRK11281 | Hypothetical protein | 0.01 | 0 | 0 | 0 | MARTX (In combination) |
| PRK12704 | PRK12704 | Phosphodiesterase | 0.01 | 0 | 0 | 0 | MARTX (In combination) |
| PRK12705 | PRK12705 | Hypothetical protein | 0.01 | 0 | 0 | 0 | MARTX (In combination) |
| PRK13914 | PRK13914 | Invasion associated secreted endopeptidase | 0.01 | 0 | 0 | 0 | MARTX (In combination) |
| PTZ00108 | PTZ00108 | DNA topoisomerase 2-like protein | 0.01 | 0 | 0 | 0 | MARTX (In combination) |
| PTZ00121 | PTZ00121 | Maebl | 0.01 | 0 | 0 | 0 | MARTX (In combination) |
| TIGR00606 | rad50 | Rad50; All proteins in this family for which functions are known are involvedin recombination, recombinational repair, and/or non-homologous end joining. | 0.01 | 0 | 0 | 0 | MARTX (In combination) |
| COG0419 | SbcC | DNA repair exonuclease sbccd ATPase subunit | 0.01 | 0 | 0 | 0 | MARTX (In combination) |
| TIGR00618 | sbcc | Exonuclease sbcc | 0.01 | 0 | 0 | 0 | MARTX (In combination) |
| pfam05483 | SCP-1 | Synaptonemal complex protein 1 (SCP-1) | 0.01 | 0 | 0 | 0 | MARTX (In combination) |
| TIGR04211 | SH3_and_anchor | SH3 domain protein; members of this protein family have a signal peptide, a strongly conserved SH3 domain, a variable region, and then a C-terminal hydrophobic transmembrane alpha helix region. | 0.01 | 0 | 0 | 0 | MARTX (In combination) |
| pfam03865 | ShlB | Haemolysin secretion/activation protein shlb/fhac/hecb | 0.01 | 0.01 | 0 | 1 | MARTX (In combination) |
| COG1196 | Smc | Chromosome segregation ATPase | 0.01 | 0 | 0 | 0 | MARTX (In combination) |
| pfam02463 | SMC_N | Recf/recn/SMC N terminal domain; this domain is found at the N | 0.01 | 0 | 0 | 0 | MARTX (In combination) |

| Accession | Short name | Definition (from InterPro database) | MARTX | RTX | YD | RTX activator | Class |
|---|---|---|---|---|---|---|---|
| | | terminus of structural maintenance of chromosomes proteins. | | | | | |
| TIGR02169 | SMC_prok_A | Chromosome segregation protein SMC | 0.01 | 0 | 0 | 0 | MARTX (In combination) |
| TIGR02168 | SMC_prok_B | Chromosome segregation protein SMC | 0.01 | 0 | 0 | 0 | MARTX (In combination) |
| pfam08317 | Spc7 | Spc7 kinetochore protein; this domain is found in cell division proteins required for kinetochore-spindle association. | 0.01 | 0 | 0 | 0 | MARTX (In combination) |
| TIGR04320 | Surf_Exclu_PgrA | SEC10/pgra surface exclusion domain. Inhibits the ability of cells to receive some plasmids | 0.01 | 0 | 0 | 0 | MARTX (In combination) |
| TIGR02680 | TIGR02680 | TIGR02680 family protein | 0.01 | 0 | 0 | 0 | MARTX (In combination) |
| COG3064 | TolA | Membrane protein involved in colicin uptake (cell wall/membrane/envelope biogenesis) | 0.01 | 0 | 0 | 0 | MARTX (In combination) |
| PRK09510 | tolA | Cell envelope integrity inner membrane protein TolA | 0.01 | 0 | 0 | 0 | MARTX (In combination) |
| TIGR02794 | tolA_full | Tola protein. The Tol-Pal complex is required for maintaining outer membrane integrity. Also involved in transport (uptake) of colicins and filamentous DNA, and implicated in pathogenesis. | 0.01 | 0 | 0 | 0 | MARTX (In combination) |
| pfam13868 | Trichoplein | Trichoplein or mitostatin, was first defined as a meiosis-specific nuclear structural protein. It has since been linked with mitochondrial movement | 0.01 | 0 | 0 | 0 | MARTX (In combination) |
| COG4717 | YhaN | Uncharacterized protein yhan, contains AAA domain | 0.01 | 0 | 0 | 0 | MARTX (In combination) |
| COG2268 | YqiK | Uncharacterized membrane protein YqiK | 0.01 | 0 | 0 | 0 | MARTX (In combination) |
| smart00191 | Int_alpha | Integrin alpha (beta-propellor repeats); integrins are cell adhesion molecules that mediate cell-extracellular matrix and cell-cell interactions | -10 | -10 | -10 | 0 | Integrin |
| cl05885 | HCBP_related superfamily | Haemolysin-type calcium binding protein related domain | 0 | 1 | 0 | 0 | Hemolysin cabind |
| pfam05860 | Haemagg_act | haemagglutination activity domain | 0.5 | 0 | 0 | 0 | MARTX |
| cl16241 | Fil_haemagg_2 | Haemagluttinin repeat | 0.5 | 0 | 0 | 0 | MARTX |

| Accession | Short name | Definition (from InterPro database) | MARTX | RTX | YD | RTX activator | Class |
|---|---|---|---|---|---|---|---|
| cl04784 | DUF637 | Possible hemagglutinin; This family represents a conserved region found in a bacterial protein which may be a hemagglutinin or hemolysin | 1 | 0 | 0 | 0 | MARTX |
| pfam09000 | Cytotoxic | The cytotoxic domain confers cytotoxic activity to proteins, enabling the formation of nucleolytic breaks in 16S ribosomal RNA | 1 | 0 | 0 | 0 | MARTX |
| cl00125 | RHOD | Rhodanese homology domain (RHOD) | -10 | -10 | -10 | 0 | Rhodanase |
| TIGR01982 | UbiB | Ubiquinone biosynthetic pathway in bacteria | -10 | -10 | -10 | 0 | Ubiquinone |
| TIGR01738 | bioH | Biosynthesis of biotin | -10 | -10 | -10 | 0 | Biotin |
